# Druggable Growth Dependencies and Tumor Evolution Analysis in Patient-Derived Organoids of Neuroendocrine Cancer

**DOI:** 10.1101/2022.10.31.514549

**Authors:** Talya L. Dayton, Nicolas Alcala, Laura Moonen, Lisanne den Hartigh, Lise Mangiante, Lisa Lap, Antonella F. M. Dost, Joep Beumer, Sonja Levy, Rachel S. van Leeuwaarde, Wenzel M. Hackeng, Kris Samsom, Catherine Voegele, Alexandra Sexton-Oates, Harry Begthel, Jeroen Korving, Lisa Hillen, Lodewijk A. A. Brosens, Sylvie Lantuejoul, Sridevi Jaksani, Niels F.M. Kok, Koen J. Hartemink, Houke M. Klomp, Inne H.M. Borel Rinkes, Anne-Marie Dingemans, Gerlof D. Valk, Menno R. Vriens, Wieneke Buikhuisen, José van den Berg, Margot Tesselaar, Jules Derks, Ernst Jan Speel, Matthieu Foll, Lynnette Fernández-Cuesta, Hans Clevers

**Author notes:** These authors contributed equally (Talya L. Dayton, Nicolas Alcala). current address: Roche Pharmaceutical Research and Early Development, Basel, Switzerland (Joep Beumer, Hans Clevers). current address: European Molecular Biology Laboratory (EMBL) Barcelona, Barcelona, Spain. Correspondence (T.L.D.), (L.F-C.), (H.C.). **In brief** Novel patient-derived tumor organoid models reveal a targetable growth-factor dependency in a subset of pulmonary neuroendocrine tumors and serve as a platform for neuroendocrine cancer research.

## Abstract

Neuroendocrine neoplasms (NENs) comprise well-differentiated neuroendocrine tumors and poorly-differentiated carcinomas. Treatment options for patients with NENs are limited, in part due to lack of accurate models. To address this need we established the first patient-derived tumor organoids (PDTOs) from pulmonary neuroendocrine tumors and derived PDTOs from an understudied NEN subtype, large cell neuroendocrine carcinoma (LCNEC). PDTOs maintain the gene expression patterns, intra-tumoral heterogeneity, and evolutionary processes of parental tumors. Through drug sensitivity analyses, we uncover therapeutic sensitivities to an inhibitor of NAD salvage biosynthesis and to an inhibitor of BCL-2. Finally, we identify a dependency on EGF in pulmonary neuroendocrine tumor PDTOs. Consistent with these findings, analysis of an independent cohort showed that approximately 50% of pulmonary neuroendocrine tumors expressed EGFR. This study identifies a potentially actionable vulnerability for a subset of NENs, and further highlights the utility of these novel PDTO models for the study of NENs.

**Graphical abstract:** 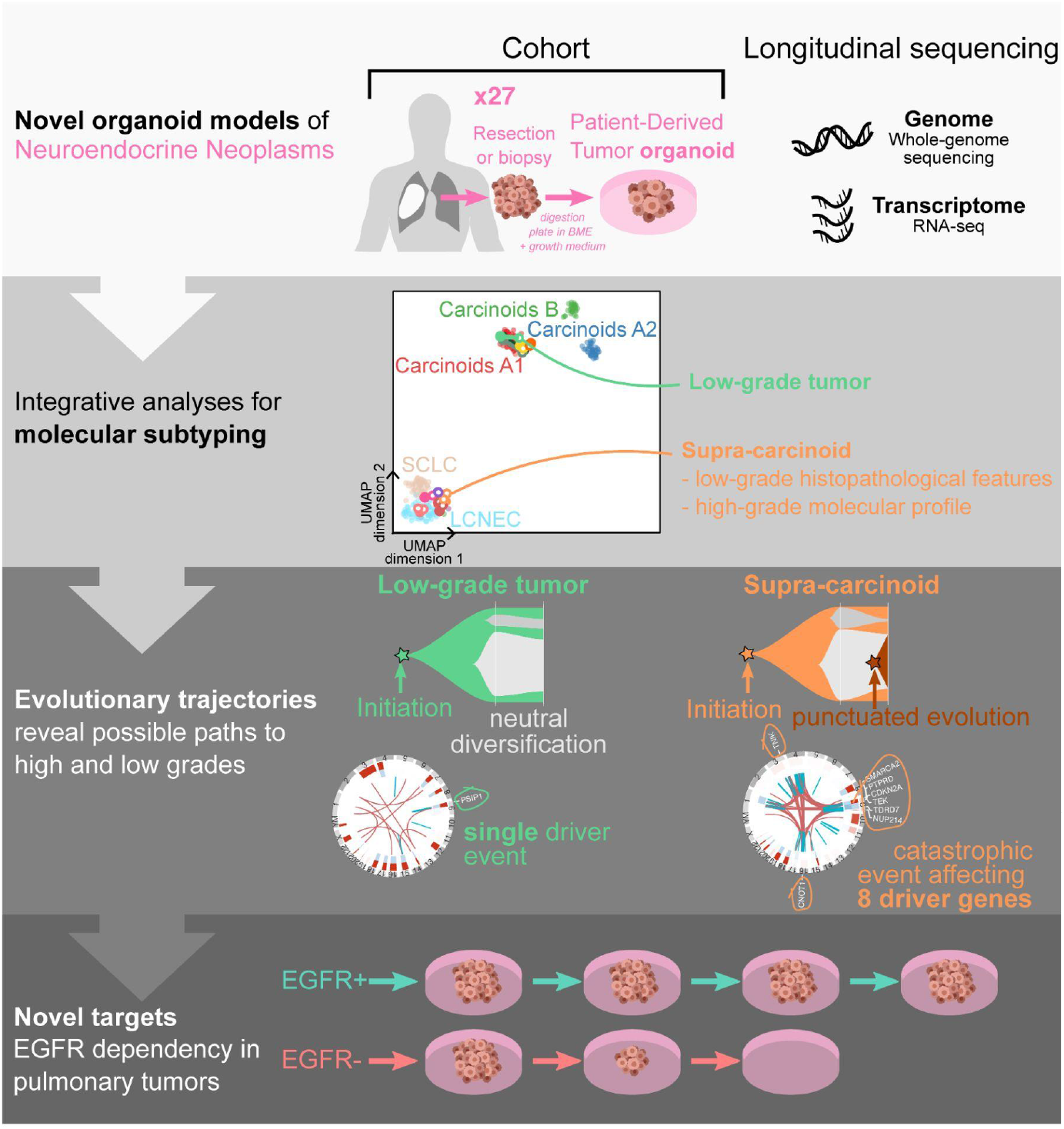

**Highlights:**

- PDTOs of pulmonary NETs and LCNEC were established
- PDTOs recapitulate intra-tumoral heterogeneity and evolution of parental tumors
- Drug assays reveal therapeutic vulnerabilities and biomarkers
- Pulmonary NET PDTOs are dependent on EGF

## INTRODUCTION

Neuroendocrine neoplasms (NENs) can arise in almost any tissue but they show the highest incidence in the lung and gastroenteropancreatic (GEP) system. These tumors show features of neuroendocrine differentiation and comprise what are considered two distinct entities, neuroendocrine tumors (NETs) and neuroendocrine carcinomas (NECs). The latter are poorly differentiated, high-grade tumors that show elevated rates of proliferation and are associated with a poor prognosis and median life expectancy of less than 1 year (Rindi et al., 2022). Based on their morphology NECs are further subdivided into small cell NECs and large cell NECs (LCNECs).

Pulmonary small cell NECs, termed small cell lung cancers (SCLCs), account for ~15% of all lung cancers, and are well-studied relative to other NEC subtypes. LCNEC on the other hand is much less well understood. Guidelines for treating patients with pulmonary and extrapulmonary LCNEC remain rudimentary. It is unclear whether the well-established therapeutic guidelines used to treat patients with SCLC can be effectively applied to patients with pulmonary LCNEC (Derks et al., 2017). Likewise, it is unknown whether pulmonary LCNEC and GEP LCNEC can be treated similarly despite arising in different tissue sites. Thus, there is an unmet need for clinical biomarkers predictive of treatment response (Baine and Rekhtman, 2020; Corbett et al., 2021).

NETs are well-differentiated neoplasms consisting predominantly of low-grade tumors (G1 and G2) that proliferate and progress slowly. The pathological WHO criteria for the diagnosis of low-grade NET is a Ki67 positivity rate of less than 20% for GEP NETs, or a maximum mitotic count of 10 mitoses per 2 mm^2^ for pulmonary NETs (Rindi et al., 2022). In the lung, low-grade NETs are referred to as typical carcinoids (TC) and atypical carcinoids (AC), corresponding to G1 and G2 tumors, respectively. Although low-grade NETs are generally associated with a favorable prognosis, up to 35% of patients with NETs present with metastases and, in these cases, overall survival rates drop significantly (Korse et al., 2013). The 10-year disease-specific survival for patients with metastatic G2 pulmonary NETs is only 18% (Baudin et al., 2021).

In the GEP system, high-grade (G3) NETs are a recognized entity defined as well-differentiated tumors that show higher Ki67 positivity rate and more aggressive clinical behavior than G1 and G2 GEP NETs. G3 NETs are associated with 5-year survival rates ranging from 7 to 40% depending on the primary site (Cives and Strosberg, 2018). Although no analogous category of G3 pulmonary NET has been officially defined, several studies have observed a subset of well-differentiated pulmonary NETs showing features of high-grade disease (Hermans et al., 2020). Two studies consisting of molecular analyses of pulmonary NET tissue showed that these tumors can be subdivided into three molecular groups: less aggressive carcinoids A1 and A2, and more aggressive carcinoids B (Alcala et al., 2019; Laddha et al., 2019). The study by Alcala et al. also identified a subgroup of NETs termed supra-carcinoids, highly aggressive tumors with the histopathological profile of low-grade NETs but the molecular profile of LCNEC. Pointing to their clinical relevance, supra-carcinoids were associated with a lower 10-year overall survival compared to G1/G2 pulmonary NETs. Other molecular analyses of pulmonary NENs have identified specific subsets of tumors with similar characteristics to supra-carcinoids (van den Broek et al., 2021; Rekhtman et al., 2016; Simbolo et al., 2019).

The lack of clarity regarding standard of care for patients with pulmonary and extrapulmonary LCNEC and unresectable, metastatic, or recurrent NETs points to two unmet needs. First, there is a need for new and effective systemic treatment strategies for these patients. Second, there is a need for suitable biomarkers to guide clinicians to choose therapeutic interventions likely to achieve the best outcomes for patients. Retrospective genomic studies have provided some insights into these NEN subtypes, generating hypotheses about potential therapeutic targets and the origin or mechanisms of progression for different molecular groups of NENs. However, to test such hypotheses, reliable preclinical models that reproduce the behavior of these different NEN entities are needed. Whereas there are several preclinical models of SCLC, including genetically engineered mouse models, more than fifty cell lines, and several tumor organoid lines, there is a relative dearth of models for pulmonary and extrapulmonary LCNEC (Gazdar et al., 2017; Kawasaki et al., 2018; Lorz et al., 2020). Similarly, only a handful of cell lines exist for NETs (Andersson-Rolf et al., 2021; Asiedu et al., 2014; Kawasaki et al., 2018; Lorz et al., 2020).

Patient-derived tumor organoids (PDTOs), which are 3D cultures of tumor cells, can be expanded long-term, can be cryopreserved, and have been shown to be representative of the patient tumor tissue from which they were derived at both the genetic and phenotypic levels (Sachs and Clevers, 2014; Sachs et al., 2018; Sato et al., 2011; van de Wetering et al., 2015). To date, a handful of PDTOs has been derived from high-grade NENs including SCLC, pulmonary LCNEC, and G3 GEP NETs and NECs (Dijkstra et al., 2021; Kawasaki et al., 2020; Kim et al., 2019; Sachs et al., 2019). To the best of our knowledge, no PDTOs of low-grade NETs have been reported. In this study, we establish a collection of PDTOs for NENs from multiple body sites, including pulmonary and extrapulmonary LCNEC and the first reported PDTOs of pulmonary NETs. We confirm the fidelity of the PDTOs at the morphological and molecular levels and show that they also preserve the intra-tumoral heterogeneity and active evolutionary processes of their parental tumors. Using this unique platform, we uncover novel therapeutic vulnerabilities for these tumors, showing their utility as a platform for neuroendocrine cancer research.

## RESULTS

### Establishment of patient-derived tumor organoids (PDTOs) of understudied NEN subtypes

To generate clinically relevant PDTO models of understudied NEN subtypes, tissue samples from patients undergoing surgical resection or biopsy for low-grade NET or LCNEC were obtained, subjected to enzymatic digestion, and the resulting cell suspensions embedded in basement membrane extract (BME) matrix and submerged in culture medium. When tumor organoids formed, they were expanded and used for downstream analyses (Figure 1A).

**Figure 1.**
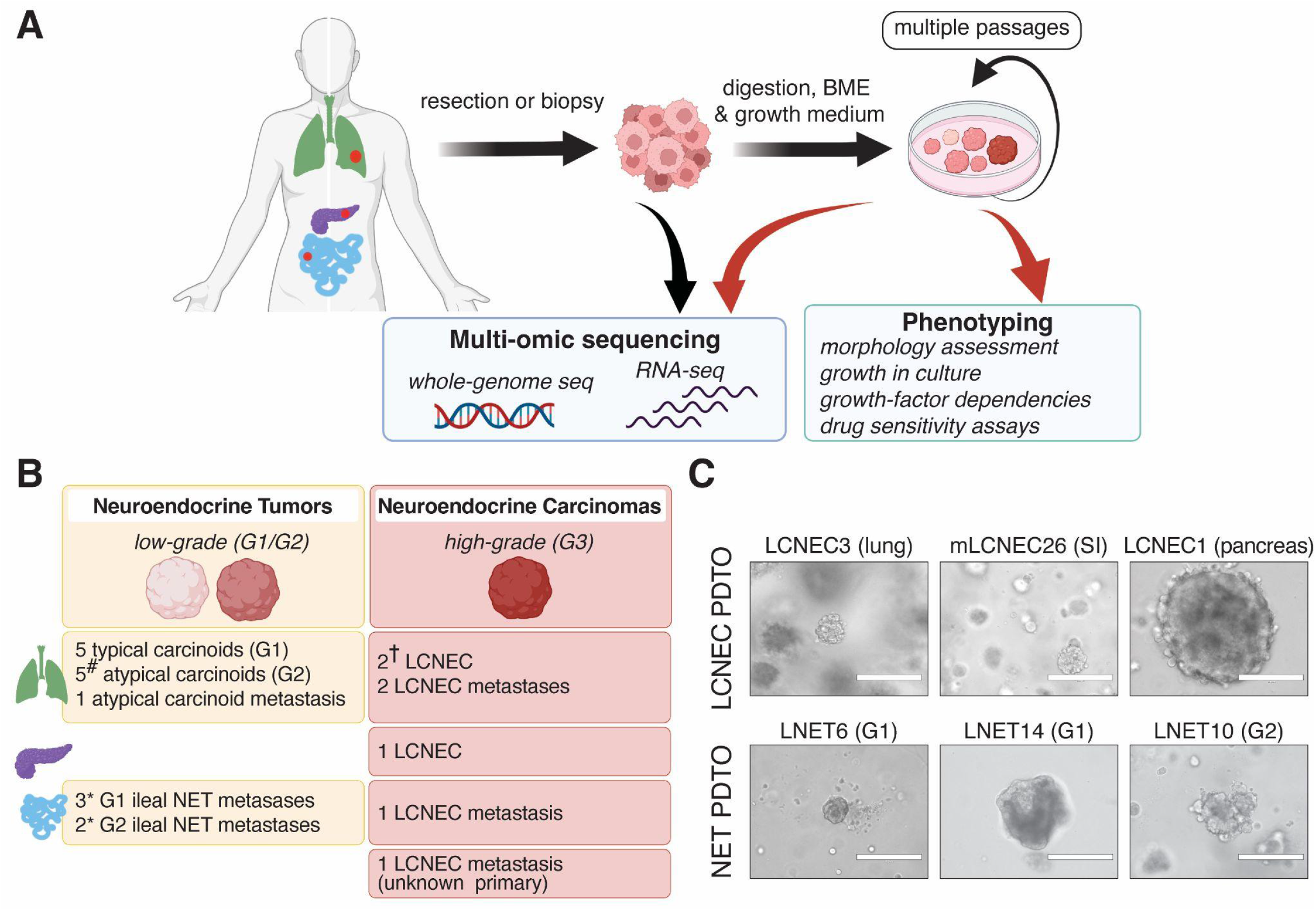
Establishment of NET and LCNEC patient-derived tumor organoids (PDTOs). (A) Schematic of experimental designs. (B) Overview of established PDTO lines. LCNEC: large cell neuroendocrine carcinoma; NET: Neuroendocrine tumor; G1: grade 1; G2: grade 2; G3: grade 3. Lung NET PDTO lines that did not show long-term growth were not included in this overview. #: 1 line from a presumed supra-carcinoid; * growth past passage 4 not achieved; †: 1 line previously reported. See also **Figure S1** and **Table S1**. A and B made with biorender.com (C) Representative bright-field images of NEN PDTOs. Scale bar: 200 μm.

A panel of 20 NET and 6 LCNEC PDTO lines was generated (Table S1). We further characterized an additional LCNEC PDTO, LCNEC4, generated and reported as part of a previous study from our lab (Sachs et al., 2019). All of the LCNEC and 11 of the NET PDTOs (all pulmonary NETs), showed long-term growth in culture (Figure 1B). One line derived from a metastasis with unknown primary, mLCNEC23, was established from a fine needle biopsy, showing the feasibility of establishing LCNEC PDTOs even from small amounts of patient material. NEN PDTOs derived from different patients showed a variety of morphologies, ranging from dense structures to grape-like cell clusters (Figures 1C and Figure S1A). Consistent with differences in their malignancy and proliferation, LCNEC PDTO lines were readily established with a success rate of 75% (6 out of 8), while pulmonary NET PDTOs showed an estimated success rate of 37% (11 out of 30; success defined as growth past 4 passages and/or 1 year) (Figure S1B).

### NEN PDTOs recapitulate disease-specific growth phenotypes

To determine whether NEN PDTOs resemble their corresponding parental tumors, whenever possible we performed histopathological analyses of PDTOs (13 out of 27 samples) (Table S1). This direct comparison showed that PDTOs captured the histological features of the matching tumor of origin (Figure 2A-C and Figure S2A-C). Immunohistochemical (IHC) staining for the neuroendocrine markers Chromogranin A (CHGA), Synaptophysin (SYP), and CD56/NCAM1 on NEN PDTOs confirmed their neuroendocrine origin (Figure 2B; Figure S2B and D). A distinguishing feature of low-grade NETs is their slow growth and low proliferation index (Rindi et al., 2022). IHC staining for the proliferation marker, Ki67 showed similar numbers of Ki67+ cells per field of view in NEN PDTOs and their matched parental tumors (Figure 2C and S2C). In line with these data, a comparison of the mRNA expression of *MKI67* in PDTOs derived from tumors of different grades confirmed that all PDTOs presented *MKI67* expression levels within what has been reported in their respective histopathological type (Alcala et al., 2019; Alvarez et al., 2018; Hofving et al., 2021) (Figure 2D).

**Figure 2.**
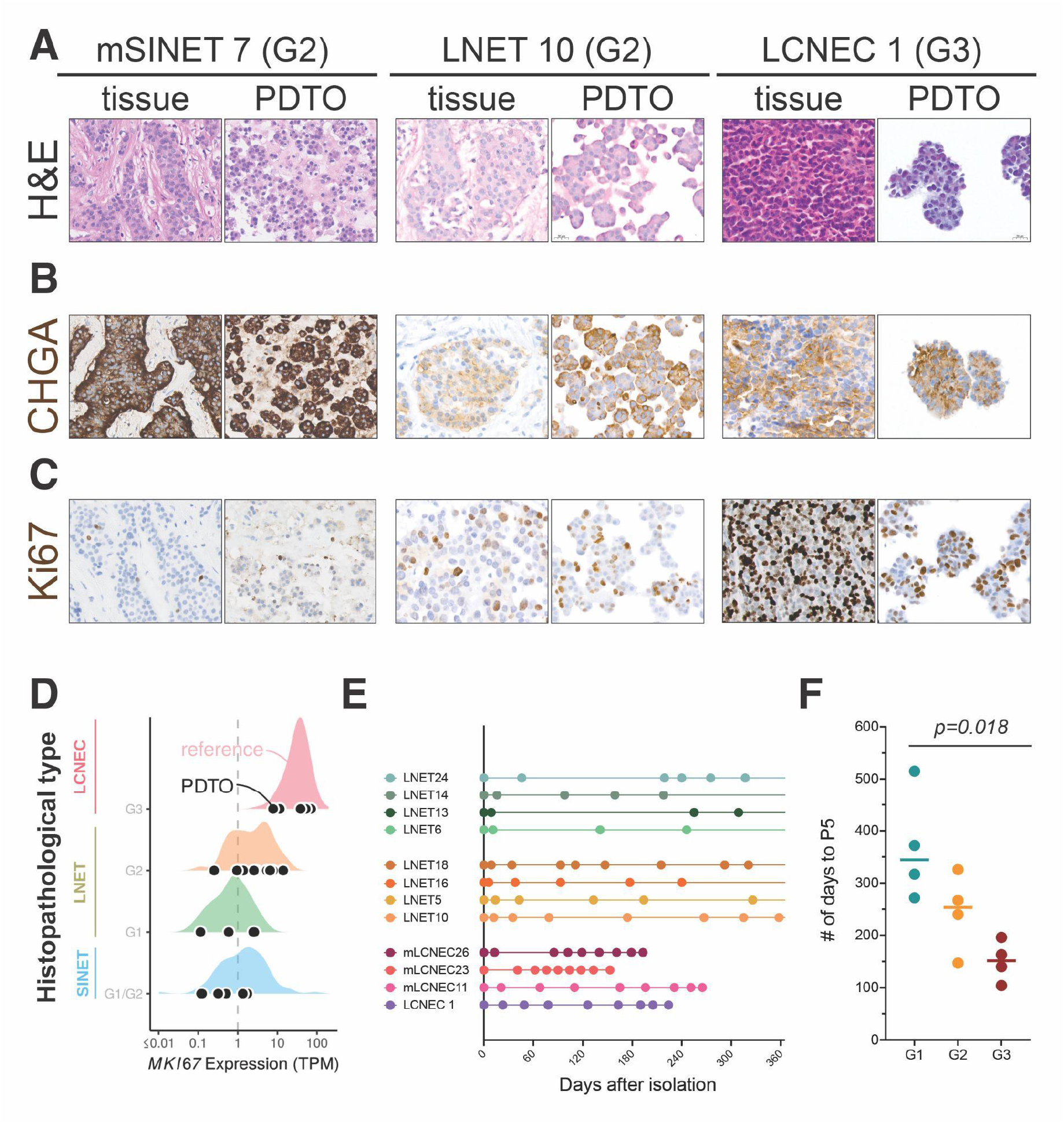
NET and LCNEC PDTOs retain histologic features and relative growth-rate of parental tumor subtypes. Representative images of (A) hematoxylin and eosin (H&E) staining and immunohistochemical staining for (B) the neuroendocrine marker, Chromogranin A (CHGA) and (C) the proliferation marker Ki67 of NEN PDTOs and their corresponding parental tumor tissue. Scale bar: 20 *μ*m. mSINET: metastasis of small intestine NET; LNET: lung NET; LCNEC: large cell NEC (D) mRNA expression of *MKI67* shown in transcripts per million (TPM) in NEN PDTO lines and reference samples of different histopathological types and grades (data from Alcala et al., 2019; Alvarez et al., 2018; Hofving et al., 2021). Black dots: PDTOs. Colored densities: distribution of expression values of reference samples. (E) Number of days in between each passage over the course of one year following isolation shown for different LCNEC and LNET PDTO lines. Each dot represents a passage. Data shown up to passage 8 or current passage number if lower than 8. mLCNEC: metastasis of LCNEC. (F) Average cumulative number of days between date of isolation and passage 5 for PDTO lines from tumors of different grades; *p* = 0.018, ANOVA See also **Figure S2** and **Table S1**.

Consistent with these observations, LCNEC and NET PDTOs had distinct *in vitro* growth rates. The average number of days it took for PDTO lines from tumors of different grades (G1, G2, or G3/LCNEC) to be passaged five times (P5) revealed a clear pattern of decreased average time to P5 with increasing tumor grade (Figure 2E and F). We did not observe a temporal trend towards deceleration or acceleration of passage time in PDTO lines (Mann-Kendall test *q*-values>0.18, Table S1). Altogether, these data argue that growth in culture did not alter tumor growth rate, which is a NEN subtype-defining phenotype.

### High-purity NEN PDTOs mirror the gene expression of their parental tumors

We next assessed whether NEN PDTOs maintained the gene expression profiles of their corresponding parental tumors using bulk RNA sequencing (RNA-seq) (Figures 3A and S3A, Table S2). To determine whether NEN PDTO samples and parental tumors show features of neuroendocrine identity, we examined the levels of expression for 3 canonical markers of neuroendocrine differentiation (*CHGA*, *NCAM1*, and *SYP*), and 3 neuroendocrine lineage transcription factors (*ASCL1*, *INSM1*, and *NEUROD1*) (Andersson-Rolf et al., 2021) (Figure 3B and S3B). For 15 out of the 21 parental tumor-PDTO families in our dataset, expression levels for these markers were similar in PDTOs and their matched parental tumors and were consistent with the range of expression observed in reference samples (Figure 3B). NET PDTOs also had expression levels of the common therapeutic target, somatostatin receptor 2 (SSTR2), consistent with the levels observed in their parental tumors (Figure S3C).

**Figure 3.**
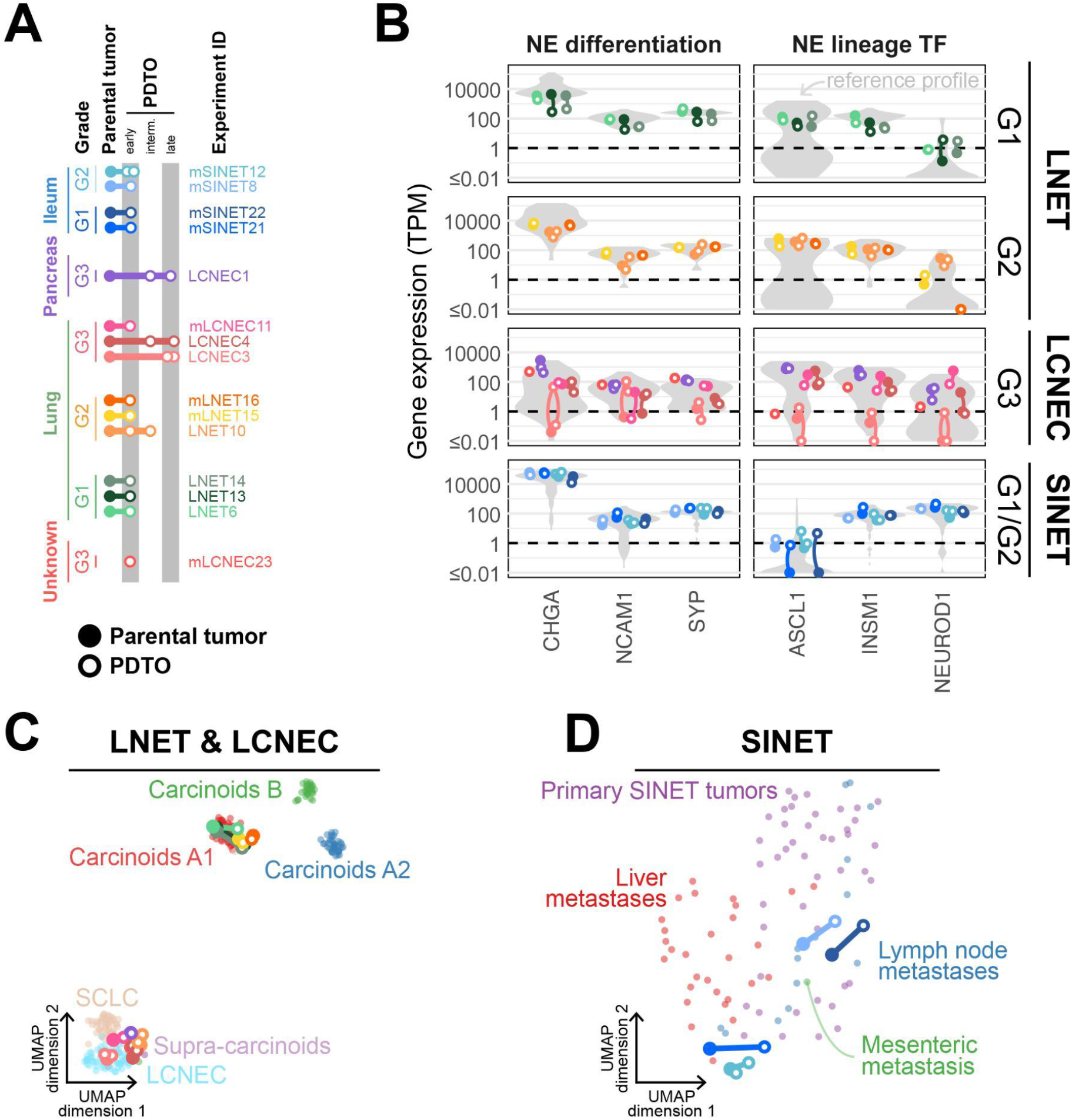
High-purity NEN PDTOs recapitulate the gene expression of original tumors. (A) Overview of high-purity PDTO lines and parental tumors for which RNA-seq data was generated. Filled circles: parental tumors; empty circles: PDTOs (early passage: 1-3; intermediate passage: 4-7; late passage: 8+). (B) Expression of neuroendocrine differentiation and transcription factor markers in parental tumors and matched PDTOs, in units of transcripts per million (TPM), for different histological types and grades. Gray violin plots represent reference profiles with matching histological type and grade (*n*=75 for G1 LNET, *n*=40 for G2 LNET, *n*=69 for LCNEC, *n*=88 for SINET). (C-D) Two-dimensional representation of the molecular profiles of PDTO within reference profiles from different molecular groups (point colors; *n*=239 LNET & LCNEC, 88 SINET) using UMAP. (C) LCNEC of the lung and pancreas for a set of 1055 genes (core genes identified in Alcala, Leblay, Gabriel, *et al*. 2019);(D) small intestine tumors for a set of 519 genes (master regulators identified in Alvarez *et al*. Nature Genetics 2018). See also **Figure S3** and **Table S2**.

The remaining 6 parental tumor-PDTO families, all derived from G1 or G2 tumors, showed levels of expression for the examined neuroendocrine markers that were lower in the PDTO than in the parental tumor tissue (Figure S3B). We performed IHC staining for the neuroendocrine marker CHGA in 3 of these PDTO lines and found that they displayed a staining pattern consistent with growth of CHGA+ tumor cells together with undefined normal cell types (Figure S3D). We, therefore, hypothesized that some PDTO lines consist of both healthy and tumor cells, while others consist primarily of tumor cells. To assess tumor purity of PDTO lines we devised a classification criterion based on a mixture of molecular and IHC criteria to classify NEN PDTOs as either “mixed” or “high-purity” (See Methods; Figure S3D; Table S3).

Transcriptomic analysis of these high-purity NEN PDTO lines and their matched parental tumors, and a comparison with transcriptomes of NEN reference samples from previously published datasets, showed that they captured the molecular group of their parental tumor (Alcala et al., 2019; Alvarez et al., 2018; Gabriel et al., 2020) (Figure 3C-D). Embedding and clustering by UMAP using genes representative of known molecular groups of pulmonary NENs and small intestinal NETs largely showed that PDTOs and their parental tumors clustered together with the reference NEN tissue samples of the expected molecular group. Although LCNEC1 and mLCNEC23 were from extrapulmonary LCNEC tumors, their matched parental tumors and PDTOs clustered with the reference pulmonary LCNECs, suggesting that LCNECs arising in different tissue sites share similar expression profiles.

The parental tumor and PDTOs from the pulmonary NET labeled LNET10 clustered with LCNEC samples. An additional three pulmonary NET samples, all from the reference datasets and previously reported as supra-carcinoids, also fell into the LCNEC cluster (Alcala et al., 2019) (Figure 3C). Consistent with the hypothesis that LNET10 is a supra-carcinoid, the parental tumor and PDTOs displayed features previously observed in supra-carcinoids: high expression levels of immune checkpoint genes and low expression of the putative prognostic marker for pulmonary NETs, *OTP* (Alcala et al., 2019; Moonen et al., 2019, 2022) (Figure S3E). The clinical data of the patient with LNET10 was also consistent with the diagnosis of supra-carcinoid: multiple metastases and recurrence following targeted therapy.

To identify markers that distinguish PDTOs from their parental tumors, we performed Partial Least Squares (PLS) analyses of all PDTO lines and their matched parental tumor (Figure S3F). We then used gene set enrichment analysis (GSEA) on the identified markers and found that they are mostly immune-related, consistent with observations in PDTOs of other tumor types (Lee et al., 2018) (Figure S3G). These data support the notion that PDTOs contain only the tumoral and epithelial components but not the stromal components of the parental tumors, such as the infiltrating or resident immune cells.

### NEN PDTOs retain the genomic profile of their parental tumors

To determine whether NEN PDTOs recapitulate the genomic landscape of their corresponding parental tumors, we performed whole genome sequencing (WGS) in 10 parental tumor-PDTO families representing all tumor types and grades in the biobank, and including multiple time points in culture for 2 LCNEC lines (Figure 4A and S4A, Table S4).

**Figure 4.**
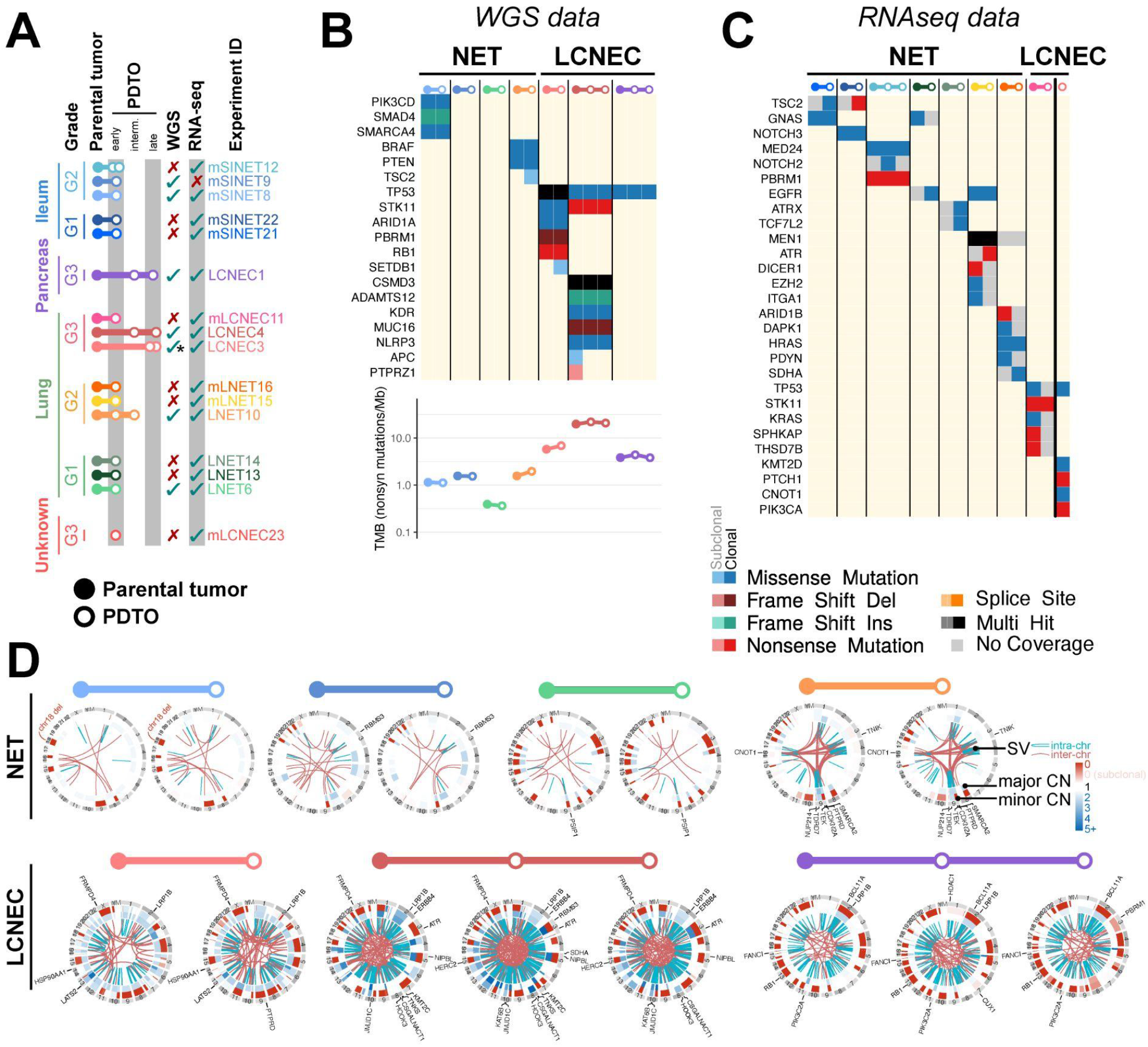
NEN PDTOs retain genomic features of parental tumors. (A) Overview of high-purity PDTO lines and parental tumors for which whole genome sequencing (WGS) and/or RNA-seq data was generated. Filled circles: parental tumors; empty circles: PDTOs (early passage: 1-3; intermediate passage: 4-7; late passage: 8+). *LCNEC3 WGS data only generated for 1 passage of LCNEC3. RNAseq data was generated for 2 passages. (B-C) Summary of damaging somatic alterations detected by (B) whole-genome sequencing or (C) RNA-seq in genes that have been reported to be recurrently mutated in LCNEC, lung NET, and SINET. Colors represent variant classes and clonality (light: subclonal; solid: clonal). In (B), the lower panel represents the Tumor Mutational Burden (TMB), expressed as the number of nonsynonymous mutations per megabase.In (C), light gray represents genes without enough coverage to detect variants. (D) Structural variants in PDTOs and parental tumors. Inner layer: chromosomal rearrangements; central layer: major copy number (CN); outer layer: minor CN. Structural variants damaging genes that have been previously reported as recurrently mutated in LCNEC, lung NET, or SINET are annotated in black. Subclonal CN alterations (non-integer CN) are indicated with intermediate colors (e.g., light red for subclonal CN loss). See also **Figure S4**, and **Table S4**.

Tumor mutational burden was similar between PDTOs and their parental tumor (Figure 4B and S4B). Consistent with previous observations (Alcala et al., 2019; Fernandez-Cuesta et al., 2014; George et al., 2018; van Riet et al., 2021), tumor mutational burden was lower for NET than LCNEC (mean 1.26 for NET and 11.5 for LCNEC). The spectrum of mutationally altered genes we observed in the NEN tumors and PDTO samples recapitulated genomic alterations and patterns previously reported in NENs (Alcala et al., 2019; Alvarez et al., 2018; Cros et al., 2021; Derks et al., 2018; Fernandez-Cuesta et al., 2014; George et al., 2018; Miyoshi et al., 2017; Pelosi et al., 2018; Rekhtman et al., 2016; Simbolo et al., 2017). NET PDTOs presented fewer single nucleotide variants (SNVs) and insertions and deletions (indels) in known driver genes than LCNEC PDTOs (average of 1.5 *vs* 4), with SINET9 and LNET6 samples showing none (Figure 4B and S4B). High-purity PDTOs showed a high degree of concordance in putative driver alterations with their corresponding parental tumors. For 5 out of 8 high-purity PDTO WGS profiles, the SNVs in putative driver genes observed in the parental tumors were also observed in the corresponding PDTO line. In the remaining 3 high-purity PDTO lines, LNET10, LCNEC3, and LCNEC4, discordance between the PDTO and corresponding parental tumor was seen exclusively in subclonal alterations, expected to be more difficult to detect, and in only up to 2 putative driver SNVs (Figure 4B). In the case of the 2 “mixed” PDTO-parental tumor WGS profiles of lung NETs in our dataset, LNET2 and LNET5, we identified somatic driver mutations that confirm that they contain tumoral tissue (Figure S4B).

To identify somatic variants in samples for which WGS had not been performed, we called small variants in genes previously found to be altered in NENs identified in the RNA-seq reads for those genes (Figure 4C and S4C). We identified 38 unique, putatively oncogenic mutations (see methods) in either the PDTOs, parental tumors, or both (Figure S4C). The mutated genes we identified involved common NEN targets, such as *MEN1*, *ATRX*, *SMARCA4*, *ARID1B*, *TSC2*, *PIK3CA*, and *STK11* (Alcala et al., 2019; Alvarez et al., 2018; Fernandez-Cuesta et al., 2014; George et al., 2018).

In all cases, when coverage of the region of interest for the relevant mutations and samples was sufficient, the mutations identified in the tumor were also found in the corresponding PDTO (true for 9/38 mutations and for 8/11 PDTO-parental tumor pairs) (Figure 4C and S4C). We identified the oncogenic *HRAS* G13D mutation in all samples from LNET16, including the primary pulmonary NET, its matched metastasis, and the corresponding tumor- and metastasis-derived PDTOs. We also identified two distinct somatic mutations in *MEN1*, the most commonly altered gene in pulmonary NETs, in both the tumor and PDTO from mLNET15 (Alcala et al., 2019; Derks et al., 2018; Fernandez-Cuesta et al., 2014). The parental tumor sample from LCNEC11 was found to carry the oncogenic *KRAS* G12V mutation and the PDTO of mLCNEC23 a predicted oncogenic mutation in the *TP53* gene. Of note, we identified 4 putative NEN-driver mutations in the sample from the “mixed” PDTO line, LNET18, suggesting that the CHGA+ cells in this line identified by IHC, are tumor cells (Figure S3D and Figure S4C).

At the level of copy number alterations and structural variants (SVs), PDTOs captured both focal events affecting only a single gene and large-scale chromosomal aberrations (Figure 4D and S4D). As examples of the former, we observed *RBMS3* translocations in SINET9, and a *PSIP1* inversion and a large deletion in LNET6. For the latter, we observed a potential chromothripsis event in LNET10 affecting chromosomes 1, 3, 4, 9, and 16, whole-genome doubling in LCNEC4, and chromosome 18 loss–a known event observed in more than 60% of SINETs–in SINET8 (Samsom et al., 2019; Zhang et al., 2020). Altogether the patterns of copy number alterations and SVs were conserved between parental tumors and PDTOs, and were consistent with patterns previously observed in and characteristic of NENs.

### Organoids capture the intra-tumor heterogeneity and evolutionary processes of their parental tumors

To determine whether PDTOs capture the fine-scale genetic makeup of their parental tumors, we performed clonal deconvolution and evolutionary analyses. PDTOs recapitulate the ratio of clonal and subclonal alterations of each tumor type (Figure 5A and S5A-B), with NETs having predominantly subclonal alterations not common to all cells in the tumor (from ~400 to ~3200), and LCNECs having predominantly clonal alterations found in all tumor cells (from ~1500 to ~5300). Importantly, subclonal alterations were predominantly present at similar cancer cell fractions in the parental tumor and matched PDTOs, and even alterations present in only a small percentage of parental tumor cells were detected in the PDTO (Figure S5A). As expected, concordance of subclonal mutations between PDTOs and parental tumors was highest in the lower grade and early passage NETs (<11% of private alterations in mSINET8, mSINET9, and LNET6). An exception to this was LNET10, the putative supra-carcinoid, which showed more substantial subclonal differences (~40% of private alterations) between parent and PDTO.

**Figure 5.**
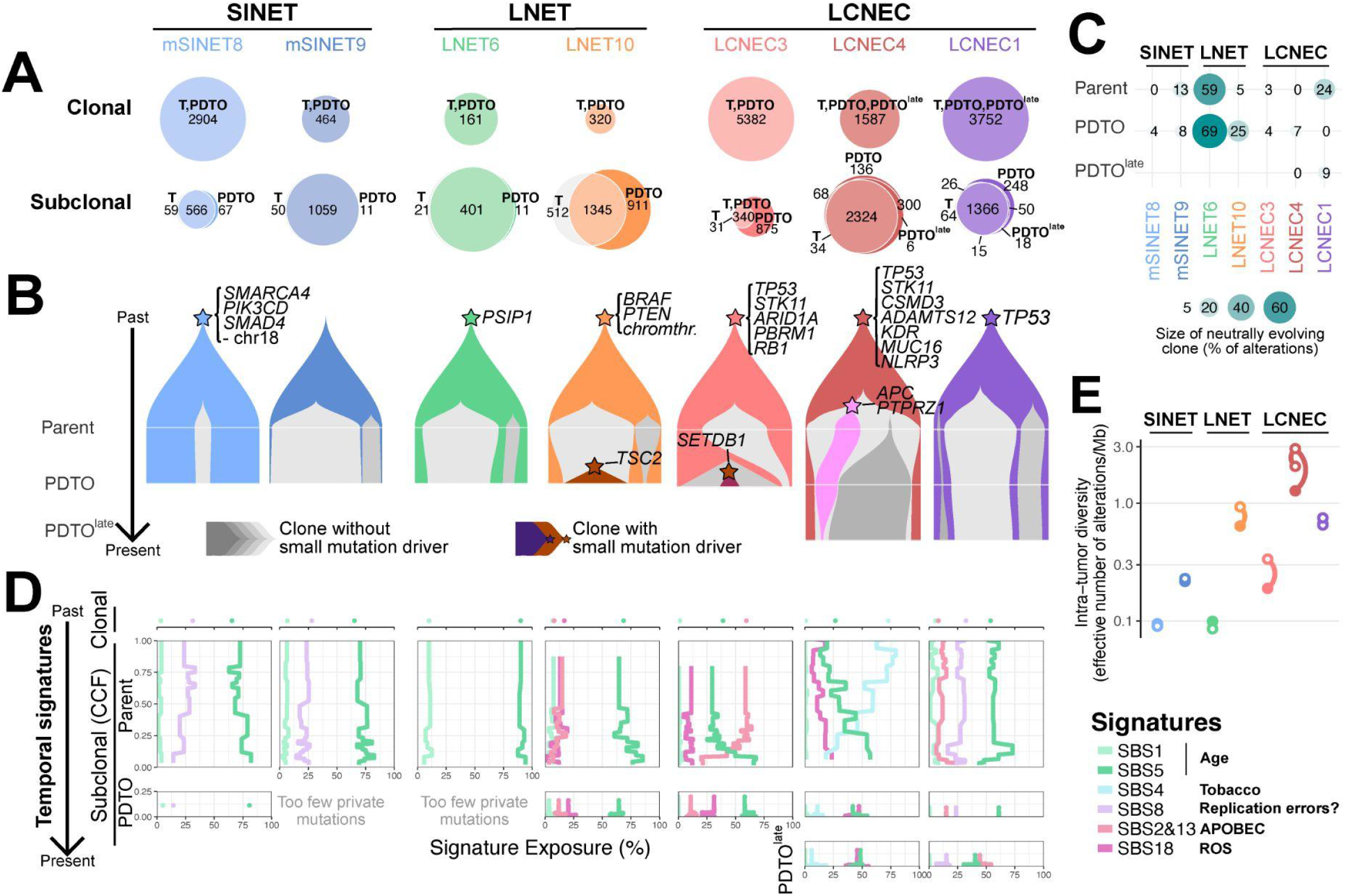
NEN PDTOs recapitulate the intra-tumor heterogeneity of the parental tumor. (A) Venn-Euler diagrams of shared and private clonal (top) and subclonal (bottom) somatic small variants. Fish plots showing clonal reconstruction of tumor and organoid. (C) Mode of evolution, measured as the size of the neutrally evolving clone in percentage of subclonal alterations (see methods). (D) Temporal mutational signatures, measured as the signature exposure (the percentage of mutations belonging to each signature), in clonal small variants (top), subclonal small variants present only in the parental tumor (middle), and those only present in the PDTO (bottom). The vertical axis corresponds to the cancer cell fraction (CCF), which is a proxy for the age of the mutation (older alterations have high CCF and are at the top, recent alterations have small CCF and are at the bottom). (E) Intra-tumor genetic diversity, measured as the effective number of alterations per Mb (see methods). In (A), (B), and– (D), columns correspond to PDTO lines/parent families. See **Figure S5** and **Table S4**.

These subclonal analyses showed that PDTOs preserve the main subclones of their parental tumor, and this was true for all samples (Figure 5B and S5C). In concordance with their lower grade and lower passage time, we did not identify known driver alterations in subclones of SINETs or LNET6 PDTOs but detected neutrally evolving subclones in both parents and PDTOs (MOBSTER method (Caravagna et al., 2020), Figure 5C and S5D), suggesting past and present neutral evolution of these tumors. Interestingly, supra-carcinoid LNET10 had multiple clonal driver mutations (*BRAF*, *PTEN*, a chromothripsis event affecting many genes), consistent with past linear evolution. This sample also had a subclonal *TSC2* mutation that was found at high frequency in the PDTO, but was either absent or below the detection limit in the parental tumor, an observation that would be consistent with a possible ongoing selective sweep. The high-grade LCNEC1, 3, and 4 also show signs of past linear evolution, and possibly ongoing natural selection, due to the presence at low frequency of driver mutations in LCNEC3 (*SETDB1*) and LCNEC4 (*APC* and *PTPRZ1*) PDTOs. These data suggest that both past and present evolution under natural selection are faithfully captured by PDTOs.

We reasoned that the mutations that were private to the PDTO could have been generated either by the same mutational processes observed in the parental tumor or by new mutational processes specific to the PDTO and/or the culture conditions. To determine which of these was true, we used the TrackSig method (Rubanova et al., 2020) to reconstruct the temporal trajectory of mutational signatures in tumors and their corresponding PDTOs.

Mutations appearing in the PDTOs were not only generated by the same processes that were active in the parental tumors, they also accurately reflected recent shifts in sources of mutation (Figure 5D and S5E). For instance, a signature associated with reactive oxygen species, single base substitution 18 (SBS18), was detected in LCNEC3 and LCNEC4 subclonal alterations and was also found in their PDTOs. Even the quasi-absence of the tobacco smoking-associated SBS4 in LCNEC4 PDTO cells is consistent with a recent drastic decline in this signature in the parental tumor (from 73% of clonal alterations to 18% of “recent” alterations, i.e., with CCF<10%). This shows that PDTOs preserve the endogenous mutational processes operating in the parental tumors.

Finally, we investigated whether PDTOs presented stable levels of intra-tumoral heterogeneity. To do so, we used the effective number of alleles to compute an effective subclonal tumor mutational burden (see Methods). Interestingly, even late passage PDTOs still harbored levels of genetic diversity similar to those observed in parental tumors (Figure 5E and S5F). No sample presented any substantial loss of diversity: LNET6, a low-grade tumor, presented the largest loss but it was very minimal (less than 0.02 effective alteration/Mb), potentially indicating that a small fraction of tumor cells did not survive in culture. These data argue that PDTOs globally preserve the evolutionary processes at work in their parental tumors: the subtle balance in parental tumors between processes generating genetic diversity, such as mutational processes, and those reducing genetic diversity, such as natural selection and genetic drift, are maintained.

### Hypothesis-driven drug sensitivity testing in PDTOs from clinically aggressive NENs

PDTOs have been shown to have a predictive value in cancer therapy (Ooft et al., 2019; Pasch et al., 2019; Sachs et al., 2018; Tiriac et al., 2018; Vlachogiannis et al., 2018; Yao et al., 2020). Given the paucity of therapeutic opportunities for patients with LCNECs, we sought to investigate whether our LCNEC PDTOs could be informative in drug sensitivity assays. We performed dose titration assays to examine the effects of several drugs on 5 LCNEC PDTOs. Given its aggressive clinical course and the molecular evidence that it represents a supra-carcinoid, we also included PDTOs derived from LNET10 in these assays.

We first tested the response of organoids to the taxane, paclitaxel, and to the mTOR inhibitor, everolimus. In the clinic, paclitaxel is used in combination with carboplatin to treat non-small cell lung cancer patients, including some patients with pulmonary LCNEC (Derks et al., 2018). Everolimus is an approved therapy for low-grade NETs also being tested in clinical trials for the treatment of patients with NECs (Andersson-Rolf et al., 2021; Christopoulos et al., 2017). We observed differential drug responses of individual PDTO lines to these compounds, with 4 out of 6 of the PDTO lines tested showing some response to both drugs (Figure 6A and S6A-B).

**Figure 6.**
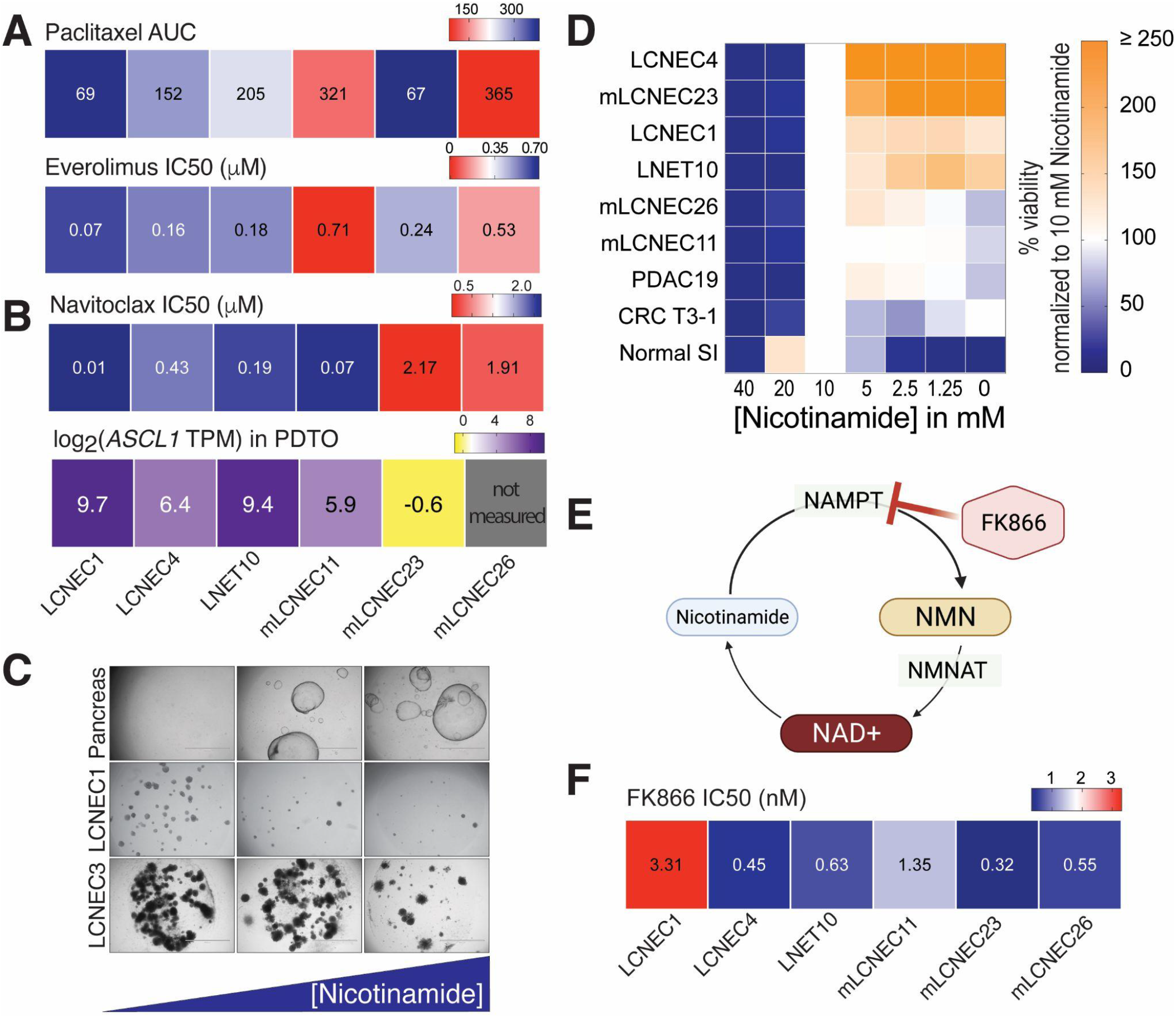
Hypothesis driven drug sensitivity testing in PDTOs from clinically aggressive NENs reveals therapeutic vulnerabilities to approved therapies. (A) Heatmap showing AUC values for Paclitaxel and IC50 values for everolimus of each NEN PDTO line tested. For Paclitaxel, AUC is reported instead of IC50 values, because the curvature of the dose response curve did not allow for IC_50_ value calculation. Dose response curves shown in Figure S6A and S6B. (B) Top: Heatmap showing IC50 values for the BCL-2 inhibitor, Navitoclax, of PDTOs. Bottom: Heatmap showing expression of *ASCL1* (in log2 of TPM) in PDTOs. (C) Brightfield images showing organoid outgrowth from single cells in media containing increasing concentrations of Nicotinamide (the main precursor for NAD+ in mammalian cells). For normal Pancreas organoids (top panel) pictures were taken on day 7 after plating. For LCNEC organoids (bottom two panels) pictures were taken on day 26 after plating. Concentrations used from left to right: 0 mM, 10 mM, 20 mM (D) Quantification of organoid outgrowth from single cells in different concentrations of Nicotinamide for LNET 10, 5 LCNEC PDTO lines, a colorectal cancer (CRC) PDTO line (T3-1), a pancreatic ductal adenocarcinoma line, normal small intestine organoids. Cell number was measured by Cell-titer Glo ATP assay and data are shown relative to outgrowth of the same line in 10 nM Nicotinamide. (E) Diagram depicting NAD+ salvage biosynthesis pathway. Created using biorender.com (F) Heat-map showing sensitivity of NEN PDTO lines to the NAMPT inhibitor, FK866, as measured by IC50 values. See **Figure S6** and **Table S5**.

We were also interested in determining whether PDTOs could predict patient response to targeted therapies. WGS analysis of the primary tumor tissue and matched PDTOs from the putative supra-carcinoid, LNET10, identified a *BRAF* V600E mutation, which leads to constitutive activation of MAPK signaling (Yaeger and Corcoran, 2019) (Figure 4B). Some patients with *BRAF*-mutated tumors show clinically significant responses to combined treatment with BRAF and MEK inhibitors (Subbiah et al., 2020). We, therefore, tested the response of LNET10 PDTOs to both single-agent and combination treatment with the BRAF inhibitor, dabrafenib and the MEK inhibitor, trametinib (Figure S6C-E). As a comparison, we tested the response of PDTOs derived from the *BRAF*-wildtype tumor (mLCNEC23) to these inhibitors. Whereas the *BRAF*-wildtype mLCNEC23 PDTOs were resistant to both single-agent treatments and their combination, the *BRAF*-mutant LNET10 PDTOs were sensitive to all treatments (AUC for combination, 186 and 96, respectively). Notably, neither drug nor their combination was able to kill all the cells in the LNET10 PDTOs. Anecdotally, treatment of the respective LNET10 patient with this same drug combination led to an initial response followed by tumor relapse and resistance to treatment.

We next sought to identify new potential therapeutic opportunities for LCNEC patients. Small cell lung cancer (SCLC) is both the most common and best studied subtype of NEC (Rindi et al., 2018). SCLC can be subdivided into molecular groups defined by the differential expression of lineage transcription factors: *ASCL1*, *NEUROD1*, and *POU2F3* (Rudin et al., 2019). Preclinical studies of SCLC suggest that tumors belonging to different SCLC molecular groups have distinct therapeutic vulnerabilities (Gay et al., 2021; Lantuejoul et al., 2020; Poirier et al., 2020; Rudin et al., 2019). *ASCL1* and *NEUROD1* are highly expressed in some LCNEC tumors, in the majority of our LCNEC PDTOs, and in LNET10 PDTOs (George et al., 2018; Hermans et al., 2019; Shida et al., 2008; Yachida et al., 2022) (Figure 3B). To determine whether these NEN PDTOs show sensitivities consistent with those identified for *ASCL1*-high or *NEUROD1*-high SCLC, we tested their response to therapies specific for these SCLC molecular groups: the BCL2 inhibitor, Navitoclax, and the Aurora kinase inhibitor, Alisertib, respectively (Gay et al., 2021) (Figure 6B and S6F). While the tested lines did not show a meaningful response to treatment with Alisertib, we noted a differential response of PDTOs to Navitoclax. In agreement with what has been observed for SCLC, PDTOs with high *ASCL1* expression were sensitive to treatment with Navitoclax, while lines with low *ASCL1* expression were resistant to the treatment. These data suggest that *ASCL1* expression in NENs might be a general biomarker for their response to BCL2 inhibitors.

Finally, we explored an observation made during the medium optimization phase of our study: NEN PDTOs showed the best outgrowth when nicotinamide was omitted from the growth medium. This was unexpected because nicotinamide is an essential component of the medium used to culture organoids derived from multiple tissue types (Huch et al., 2013; Sato et al., 2011). To quantitatively assess the effect of nicotinamide on NEN PDTOs, we tested the outgrowth efficiency of single cell suspensions derived from LCNEC and LNET10 PDTOs, from PDTOs of other tumor types (colorectal cancer and pancreatic ductal adenocarcinoma), and from healthy tissue derived organoids, in different concentrations of nicotinamide (Figure 6C and 6D). As previously observed, 10 mM nicotinamide was optimal or near optimal for outgrowth of the healthy tissue organoid lines and the colorectal cancer PDTO line. However, this was not the case for NEN PDTOs, where 5 out of 6 NEN PDTO lines showed the best outgrowth at either the lowest concentration of nicotinamide, 1.25 mM, or in its complete absence.

Nicotinamide is a precursor of oxidized nicotinamide adenine dinucleotide (NAD+), which influences energy metabolism and cellular redox states (Verdin, 2015). We speculated that the outgrowth inhibition exerted by nicotinamide on NEN PDTOs might implicate a sensitivity of NEN tumor cells to drugs that influence the conversion of nicotinamide to NAD+ in the cell. To test this possibility, we exposed LCNEC PDTOs and LNET10 PDTOs to FK866, an inhibitor of nicotinamide phosphoribosyltransferase (NAMPT), the enzyme that catalyzes the rate-limiting step in the NAD+ salvage pathway that generates NAD+ from nicotinamide (Verdin, 2015) (Figure 6E-F and S6G). Although the tested NEN PDTO lines showed individual differential responses to treatment with FK866, all the tested lines were sensitive to the treatment, and 5 out of 6 PDTO lines showed subnanomolar IC_50_ values. These results indicate that the media requirements of PDTOs can be useful for identification of potential therapeutic vulnerabilities.

### A subset of lung NET PDTOs are dependent on EGF and expression of the EGF receptor is common in lung NETs

To identify dependencies in NEN PDTOs, we tested the requirement for growth factors commonly used in organoid culture: EGF or FGF7 and FGF10. When sufficient tissue was available at isolation, we cultured a portion of the cell suspension in base NEN PDTO media, a portion in media supplemented with FGF7 and FGF10, and a portion in media supplemented with EGF. Consistent with the notion that high-grade tumors acquire growth-factor independence, 5 out of 7 LCNEC PDTOs showed no discernible difference in growth with the additional growth factors (Figure S7A).

Pulmonary NET PDTOs appeared to be EGF-dependent. Starting at the time of organoid isolation, we directly compared the outgrowth of five pulmonary NET PDTOs in media containing EGF and media lacking EGF (Figure 7A). In all cases, better outgrowth was observed in media containing EGF. In 4 out of the 5 cases, the omission of EGF precluded growth past 3 passages. SINETs did not show growth beyond passage 4 and were therefore not tested for growth factor dependencies.

**Figure 7.**
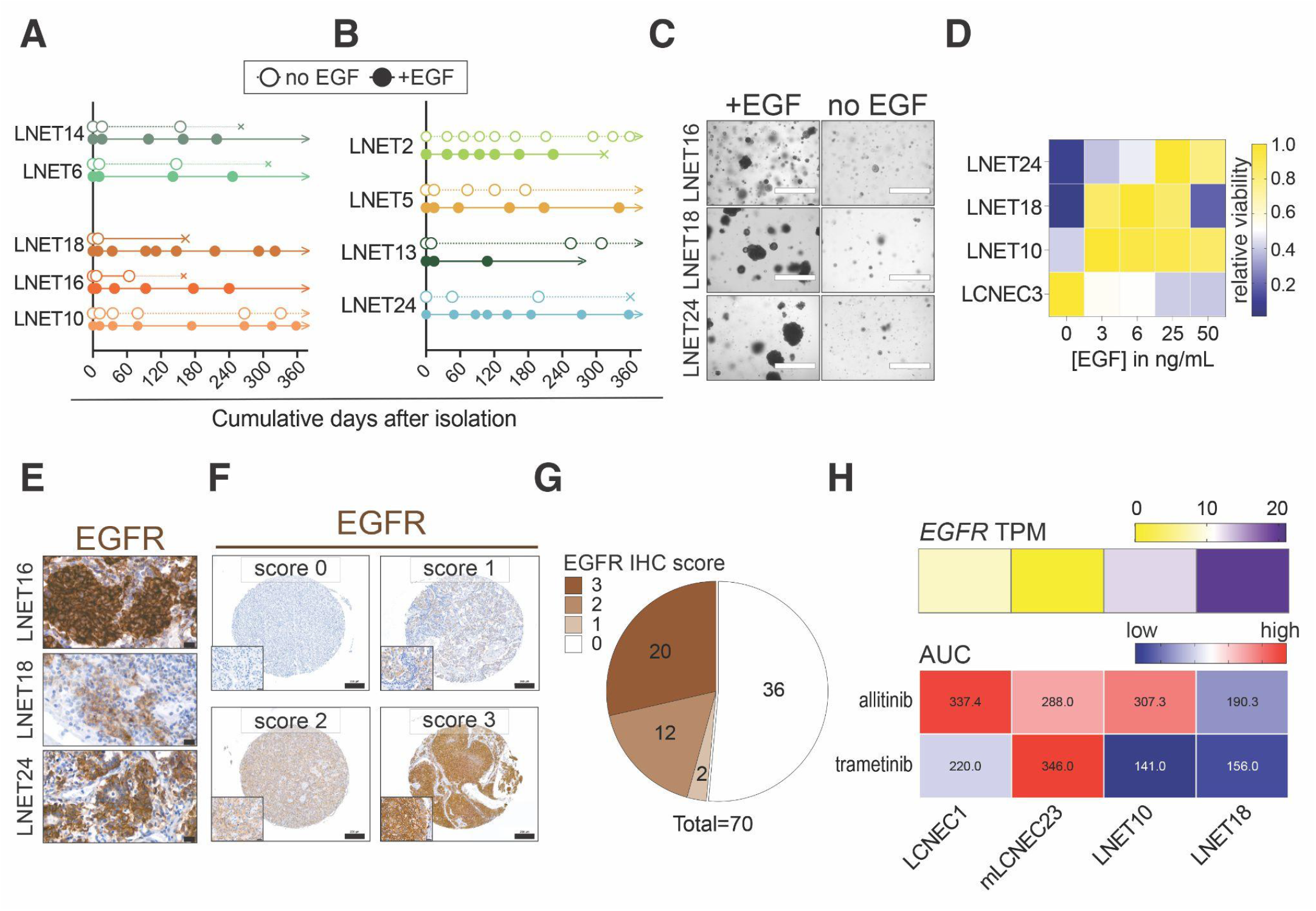
Lung NET PDTOs are dependent on EGF and EGFR expression is common in lung NETs. Number of days in between each passage over the course of one year (A) following isolation in medium with or without EGF, or (B) following isolation into medium without EGF or thawing of early passage organoids into medium with EGF. Each dot represents a passage. (C) Brightfield images showing growth of lung NET PDTOs in the presence or absence of EGF, 31 days (LNET16), 42 days (LNET18), or 75 days (LNET24) after plating. Quantification of organoid outgrowth from single cells in different concentrations of EGF for 3 pulmonary NET PDTOs and 1 LCNEC PDTO. Cell number was measured by Cell-titer Glo ATP assay and data are shown relative to outgrowth of the same line in no EGF and normalized to highest viability value. Immunohistochemical staining for the EGF receptor, EGFR, in parental tumor tissue for the corresponding PDTO lines shown in C. Scale bar: 20 μm (F) Representative tissue microarray (TMA) cores containing pulmonary NET samples stained for EGFR. An example of each EGFR IHC intensity score is shown (0, negative; 1, weak; 2, medium; 3, strong). Intensity scores were determined by percent of positive cells and intensity of membrane and cytoplasmic staining. See **Table S5** for a summary of scores and percent positive tumor cells for each TMA core and tumor. Scale bars: 200 μm; inset: 20 μm (G) Distribution of EGFR IHC intensity scores for 70 lung NETs from two different TMAs. To capture potential heterogeneity across tumors, each tumor in the TMA was represented by 3 cores. (H) Heat-map showing sensitivity of LNET 18, LNET 10, and LCNEC PDTO lines to the EGFR inhibitor allitinib, and the MEK inhibitor, trametinib, as measured by area under the curve (AUC). Numerical values for AUC are shown. Red indicates high AUC values, blue indicates low AUC values. AUC was used because the curvature of the kill curve did not allow for IC _50_ value calculations. Expression of *EGFR* for each PDTO line tested are also shown (in TPM). See **Figure S7** and **Table S6**.

Given that pulmonary NET PDTOs were dependent on EGF, we asked whether other LNET PDTO lines that had shown suboptimal growth in medium without EGF, might be similarly dependent on EGF. We then thawed frozen vials of 4 pulmonary NET PDTO lines (LNET2, LNET5, LNET13, and LNET24), directly into media containing EGF and their outgrowth over time was compared to the outgrowth dynamics we had observed for the same lines in media without EGF (Figure 7B). Indeed, 2 out of these 4 PDTO lines (LNET5 and LNET24), could be expanded more times within a 1-year time frame than when they had been grown for the same length of time without EGF. LNET24 displayed the most striking difference; without EGF it could only be passaged 3 times before being discarded due to negligible growth, but with the addition of EGF this PDTO could be passaged 7 times. In line with these observations, withdrawal of EGF from lines grown in EGF media, or addition of EGF to lines that had been grown in the absence of EGF, revealed improved expansion in the EGF-containing media (Figure 7C).

To quantitatively assess EGF-dependency in LNET PDTOs, we performed an EGF dose response outgrowth assay (Figure 7D). Consistent with our previous results, EGF improved the expansion of LNET18 and LNET24 PDTOs in a dose dependent manner up to a final EGF concentration of 25 ng/μL. While 2 out of 3 LNET PDTOs showed similar expansion at 25 ng/μL and 50 ng/μL, LNET18 PDTOs showed reduced expansion at 50 ng/μL of EGF compared to even 3 ng/μL of EGF. LCNEC3 PDTOs were not dependent on EGF and we noted the best expansion in the absence of EGF. Given that LNET10 tumor tissue and PDTOs harbor a *BRAF^V600E^* mutation, which leads to EGF-independent activation of the MAPK pathway, we were surprised to observe that LNET10 PDTOs expanded more in all of the tested EGF concentrations compared to in media where EGF was omitted. Adaptive feedback activation of MAPK signaling has been observed in *BRAF*-mutant colon cancers treated with BRAF inhibitors, suggesting that EGF-mediated MAPK activation can play a functional role in promoting the growth of EGFR-expressing *BRAF*-mutant tumors (Capdevila et al., 2020; Prahallad et al., 2012).

We next asked whether LNET primary tumor tissue and matched PDTOs expressed the EGF receptor, EGFR. For this purpose, we analyzed RNAseq data from PDTOs and parental tumors and from previously published datasets of pulmonary NET tissue (Alcala et al., 2019) (Figure S7B). *EGFR* expression was present in most tumors and PDTOs in these datasets. IHC staining for EGFR confirmed membrane EGFR protein expression in tumor cells of 11 out of 13 LNET parental tumors (Figure 7E and S7C, Table S5). In line with the observation that LNET2 PDTOs were not dependent on EGF and that LNET19 PDTOs could not be propagated in EGF-containing media after passage 3, the staining for EGFR on LNET2 and LNET19 parental tumor tissue was entirely negative. LNET15 and LNET20 parental tumor tissue showed some staining for EGFR but their PDTOs stopped expanding at passage 3, suggesting that other factors beyond EGF might contribute to *in vitro* growth of some EGFR-expressing tumors.

To ask whether EGFR expression is a common feature of pulmonary NETs, we performed IHC staining for EGFR on two sets of pulmonary NET tumor tissue microarrays containing a total of 216 cores from 73 pulmonary NETs (Figure 7F). We assigned each stained core a membrane EGFR staining intensity score, and used this to assign a score to the pulmonary NETs on the array. 48% of tumors expressed membrane EGFR, with the majority of these showing very strong staining (Figure 7G). Altogether, these data show that a subset consisting of close to half of pulmonary NETs expresses membrane EGFR.

The dependence of LNET PDTOs on EGF and the observation that a subset of pulmonary NETs express EGFR imply that some pulmonary NETs might be sensitive to treatment with inhibitors of either EGFR or EGFR downstream signaling such as MEK inhibitors. To begin to test this possibility, we treated 2 LNET PDTO lines (LNET16 and LNET10), and 2 LCNEC PDTO lines (LCNEC1 and mLCNEC23), with the EGFR inhibitor Allitinib, and the MEK inhibitor Trametinib (Figure 7H). LNET18 PDTOs, which were the only PDTOs tested that were EGF-dependent, were also the only lines showing appreciable sensitivity to EGFR inhibition (AUC of 190 compared to >300 for all other tested lines). This line was also sensitive to treatment with the MEK inhibitor. Of note, LCNEC1 and LNET10 were also both sensitive to MEK inhibition, indicating that MAPK signaling might be important for a subset of NENs independent of tumor grade (NET vs NEC). Collectively, these data are consistent with the notion that a subset of pulmonary NETs is dependent on EGF growth-factor signaling and provide a rationale for further investigating the potential for treating these tumors with EGFR or MAPK-targeted therapies.

## DISCUSSION

In this study, we established a biobank of human NEN PDTOs that fully recapitulates the spectrum of malignancy observed for NENs, encompassing both slow growing tumors and highly proliferative and metastatic carcinomas. Our biobank includes models of an understudied subtype of high-grade NEN, LCNEC, the first described PDTOs of low-grade NETs, and a PDTO derived from a supra-carcinoid, a clinically aggressive, well-differentiated pulmonary NET. Other NEN organoid biobanks were recently reported (Dijkstra et al., 2021; Kawasaki et al., 2020). However, these biobanks contained primarily GEP NECs or GEP G3 NETs and lacked models of low-grade NETs and pulmonary LCNEC. Lung NENs account for 25% of all NENs and even low-grade pulmonary NETs show a significant rate of metastasis (Andersson-Rolf et al., 2021; Korse et al., 2013). Thus, the inclusion of low-grade pulmonary NET PDTOs in our collection represents a valuable addition to the cell models currently available for research on NENs. Altogether, our NEN PDTO biobank will provide additional opportunities for investigating carcinogenesis and therapeutics across the broad spectrum of aggressiveness for the disease.

### Experimental models for low-grade tumors

A hurdle in the development of NET PDTOs has been the lack of clarity regarding their potential growth-factor dependencies. The presumed cells of origin for NETs, neuroendocrine cells, are highly specialized differentiated cells for which proliferative signals have not been entirely delineated. Our findings relating to the EGF dependence of some pulmonary NET PDTOs have important clinical implications. First, pulmonary NETs that express membrane EGFR might be amenable to therapeutic strategies that either directly target this receptor, or that target receptor-mediated downstream signaling pathways and that are currently approved for the treatment of other tumor types. There have been reports of EGFR expression in pulmonary NETs, but these studies were unable to determine whether receptor expression played a functional role in promoting the growth or survival of these tumors (Bago-Horvath et al., 2012; Bischoff et al., 2022; Rickman et al., 2009; Rusch et al., 1996). Our data argue the expression of EGFR is indicative, to some degree, of EGF-dependence. The patient population most likely to benefit from EGFR- and EGFR pathway-targeted treatments could be identified through immunohistochemical staining of tumor tissue for EGFR. Given that not all the EGFR-expressing tumors in our study could be propagated as PDTOs despite the addition of EGF to the medium, research aimed at identifying additional biomarkers of EGF-dependence would be needed. We were only able to test the response to an EGFR inhibitor of one relevant pulmonary NET PDTO line and studies in additional PDTO lines are warranted.

A question that arises when thinking about the EGF-dependence of some pulmonary NET PDTOs, is whether this is an example of a more generalizable principle. Based on our data, we hypothesize that NETs are generally growth-factor dependent. In that case, the identification of additional, potential NET-subtype specific, growth-factor dependencies could enable the generation of PDTO models of additional NETs subtypes, such as ileal NETs, which we were not able to propagate long-term. Consistent with this idea, a recent report found that certain polymorphisms in *EGFR* were associated with increased risk of developing pancreatic NET (Marinović et al., 2022). Beyond model generation, the identification of growth-factor dependencies for these and other tumors could lead to the identification of new therapeutic strategies aimed at targeting growth-factor mediated pathways in specific patient populations.

### PDTOs allow modeling of plausible evolutionary scenarios

Our comprehensive analysis of the genomic features of NEN PDTOs highlights both the fidelity and utility of these models also for research on intra-tumoral heterogeneity dynamics and tumor evolution. We saw that PDTOs not only retain most of the intra-tumoral heterogeneity of their parental tumors, they also recapitulate the evolutionary forces at work (mutation patterns, natural selection, growth) and the dynamic nature of tumor subclones. New tumor subclones are expected to appear periodically in a tumor, and older subclones either to increase in frequency until becoming clonal, or to decrease in frequency and disappear (Bozic and Wu, 2020; Durrett et al., 2011). Recent analyses of more than 2,500 tumors showed that the age of subclones depends on the aggressiveness of the disease, and ranges from a few months in highly aggressive tumors (e.g., lung adenocarcinoma and squamous cell carcinoma) to a few years for lowly aggressive tumors (Gerstung et al., 2020). Because our experiments spanned more than a year, we thus expected to observe such turnovers, and indeed we found that the late passage G3 PDTOs (e.g., passage 14 in LCNEC1, passage 17 in LCNEC3), collected more than a year after their parental tumor counterpart, show changes in subclonal composition, while low-grade tumors show no such turnover (e.g., in passage 2 of SINET8 collected after approximately 3 months). The supra-carcinoid PDTO also showed a turnover speed similar to that of LCNEC, in line with other evidence suggesting a more aggressive disease (Alcala et al., 2019). These data argue that PDTOs can be harnessed to study subclonal tumor cell dynamics and tumor evolution. This is critical to accurately model disease treatment response, including disease resistance and relapse, which is often driven by either pre-existing, low-frequency subclones or novel subclones appearing after the onset of therapy (Marusyk et al., 2020).

### Our detailed analyses shed light on NET biology

Despite our low sample size, our molecular analyses of NEN PDTOs and their parental tumors highlight illuminating cases that provide novel findings on pulmonary NETs. We saw that evolutionary trajectories can strongly vary across LNETs of different grades and molecular groups. Low-grade tumors such as LNET6 can be initiated by as little as a single driver alteration (in this case, a *PSIP1* structural variant, a common event in LNETs (Fernandez-Cuesta et al., 2014), followed by the slow accumulation of neutral (non-driver) alterations under the influence of weak age-related endogenous mutational processes. At the other end of the spectrum, supra-carcinoids such as LNET10 can evolve following catastrophic chromosomal events such as chromothripsis, that simultaneously affect multiple cancer genes, fueled by more diverse mutational processes spanning small variants and large structural rearrangements. This is a textbook example of the punctuated evolution model, whereby tumor evolution can be stagnant for a few years before undergoing a “leap” due to a sudden catastrophic event (Gould and Eldredge, 1972; Vendramin et al., 2021). Interestingly, this supra-carcinoid also seems to experience an additional subsequent selective sweep and to undergo a fast allelic turnover. A recent report from the Dutch MEN1 Study Group presented a case report where, consistent with a model of progression via punctuated evolution, a pulmonary NET showing an indolent course for six years unexpectedly changed course and progressed to aggressive disease likely driven by an activating mutation in *PIK3CA* (van den Broek et al., 2021). Further studies will be needed to determine whether these observations can be generalized and to what degree they represent a consistent model of how slow growing tumors can progress to become clinically aggressive.

### Drug screens in PDTOs of aggressive NEN subtypes highlight their utility as preclinical models

PDTOs enable studies aimed at identifying both novel therapeutic strategies and biomarkers of treatment response. Here we report case studies in drug response in LCNEC and supra-carcinoid PDTOs that highlight their utility in this regard. Consistent with studies showing that solid tumors with a more neuroendocrine phenotype show increased sensitivity to an inhibitor of the NAD salvage pathway, FK866, we found that NEN PDTOs are sensitive to this inhibitor (Balanis et al., 2019). It is tempting to speculate that the neuroendocrine phenotype, associated with the biosynthesis of secreted hormones and neuropeptides, might create specific metabolic vulnerabilities that could be exploited therapeutically.

We also found that high expression of *ASCL1* might serve as a biomarker for therapeutic response to the BCL2 inhibitor, Navitoclax. Although *ASCL1* has been identified as a biomarker of response to BCL2 inhibitors in SCLC, this therapy and the link to *ASCL1* expression has not been explored in LCNEC. The fact that we did not see the same link between *NEUROD1* expression and sensitivity to Aurora kinase inhibition that has been observed for SCLC argues that not all SCLC therapies and biomarkers can be translated to LCNEC, and underscores the need for more preclinical *in vitro* models of LCNEC where such hypotheses can be tested. While *ASCL1* has been more classically associated with pulmonary NENs, its overexpression has been recently reported also in some GEP and prostate NECs and our data show that expression of this neuroendocrine transcription factor may have clinical relevance for both pulmonary and extrapulmonary LCNECs (Cejas et al., 2021; Kawasaki et al., 2020; Yachida et al., 2022). The question that follows then is, to what degree are LCNECs from different tissue sites similar and could therapeutic strategies and biomarkers of response be applied across tumors of different tissue sites? The LCNEC PDTO samples in this study are not enough to make such a broad generalization, but our data support the idea, and further research is warranted. Indeed, the ability to classify LCNECs according to shared gene expression or drug sensitivity profiles, irrespective of tissue site, could aid in overcoming some of the obstacles associated with the fact that LCNEC, when divided according to tissue site, is very rare at each site.

## Conclusion

In conclusion, analysis of the PDTOs in our NEN PDTO library with regards to their unique media dependencies and their drug response, combined with a comprehensive examination of their genomic features allowed us to derive insights into the biology of NENs. We identified potentially actionable vulnerability for both low-grade and high-grade disease, highlighting the importance of preclinical models for the entire spectrum of malignancy encompassed by NENs. It will be important to follow up on our findings and to increase the size of this biobank to capture other NEN subtypes. Our pulmonary NET PDTOs, the first thus far reported, represent an important resource for the study of this enigmatic disease and will open the door to studies aimed at identifying the mechanisms by which they progress to stage IV disease, and the factors that predict the likelihood of this progression for a given tumor.

## ACKNOWLEDGEMENTS

We thank all patients participating in this study as well as the teams approaching patients for consent and collecting tissue, including the Utrecht Portal for Organoid Technology (U-PORT; UMC Utrecht) – in particular Anneta Brousali, Jorieke Salij, Onno Kranenburg, and Renate Bezemer – and the clinical studies department at the NKI – in particular Jan-Nico Ridderbos. We are grateful to the Foundation Hubrecht Organoid Technology (HUB) and employees, including Patrick de Kort and Calinda Dingenouts, for their work supporting ethical regulatory affairs. We thank Utrecht Sequencing for bulk RNA-sequencing services. We acknowledge financial support from the NET Research Foundation (2017 Petersen Accelerator Award to H.C.), Worldwide Cancer Research (2020 Grant Round to L.F-C), NET Research Foundation (2019 Investigator Award to L.F-C), French National Cancer Institute (INCa, PRT-K 2017 to L.F-C. and M.F.), Ligue Nationale contre le Cancer (fellowship to L.Ma.), and the Dutch Cancer Foundation (grant number 10956, 2017, to E.J.S.). T.L.D. was supported by an EMBO long-term fellowship (ALTF-21-2017) and a Marie Skłodowska-Curie IF grant 797966 – PNECtumor. A.D. was supported by Accelerate Lung Regeneration Consortium grant BREATH 12.0.18.002 of the Lung Foundation Netherlands (to H.C.). The Oncode Institute is supported by the Dutch Cancer Society.

The results shown here are in part based upon data generated by the Rare Cancers Genomics initiative (www.rarecancersgenomics.com) and the TCGA Research Network (https://www.cancer.gov/tcga).

We thank E. Reckzeh and Y.M. Soto-Feliciano for critically reading the manuscript and R. Millen for assistance with drug sensitivity assays.

## AUTHOR CONTRIBUTIONS

T.L.D., N.A., L.F-C., and H.C. designed and conceived the study. T.L.D. generated the organoids. T.L.D. and L.dH. cultured and curated organoid lines and performed related experiments. T.L.D. and A.F.M.D. performed drug sensitivity assays. J.B. and J.D. contributed to study design. L.Mo., L.L., W.M.H. and L.B. performed histological stains, acquired images, and performed related analyses. N.A. processed the sequencing data. T.L.D. and N.A. analyzed and interpreted the data and generated figures. J.V. and H.B. embedded organoids and cut slides for immunohistochemistry staining. S.J. generated novel reagents/model. L.H., L.B., and S.L. performed pathological assessments. N.A., C.V. and A.vH. curated sequencing data. L.Ma. contributed to analysis of sequencing data. L.Mo., J.D., S.L., R.S.vL., L.B., K.S. provided samples, curated patient data, and provided pathology information. N.F.M.K., K.J.H., H.M.K., I.H.M.B.R., A-M.D., G.V., M.R.V., W.B., J.vdB., M.T. provided clinical feedback. J.D., M.R.V., G.D.V., J.vdB., and M.T. coordinated clinical aspects of the study. L.F-C. and M.F. supervised N.A., L.Ma., C.V., and A.S-O. E-J.S. and J.D. supervised L.Mo. and L.L. H.C. directed the study.

## METHODS

### KEY RESOURCES TABLE

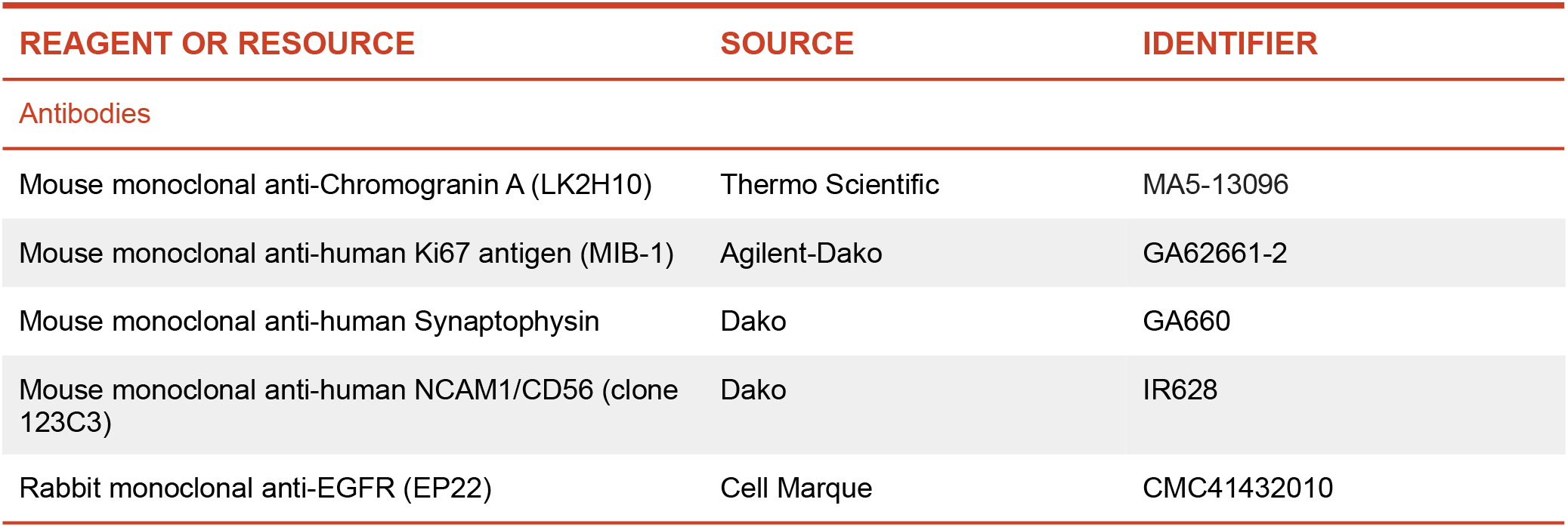
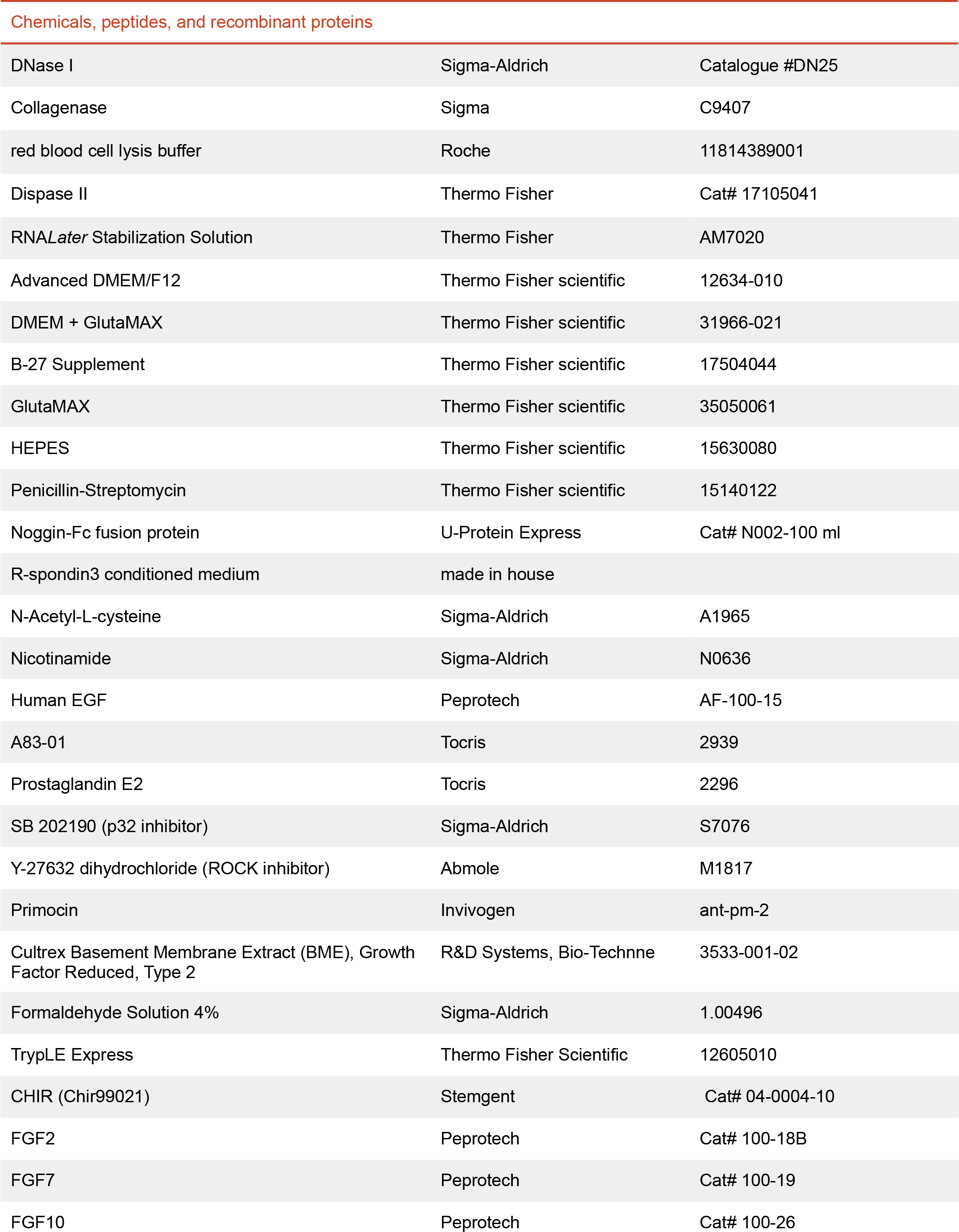
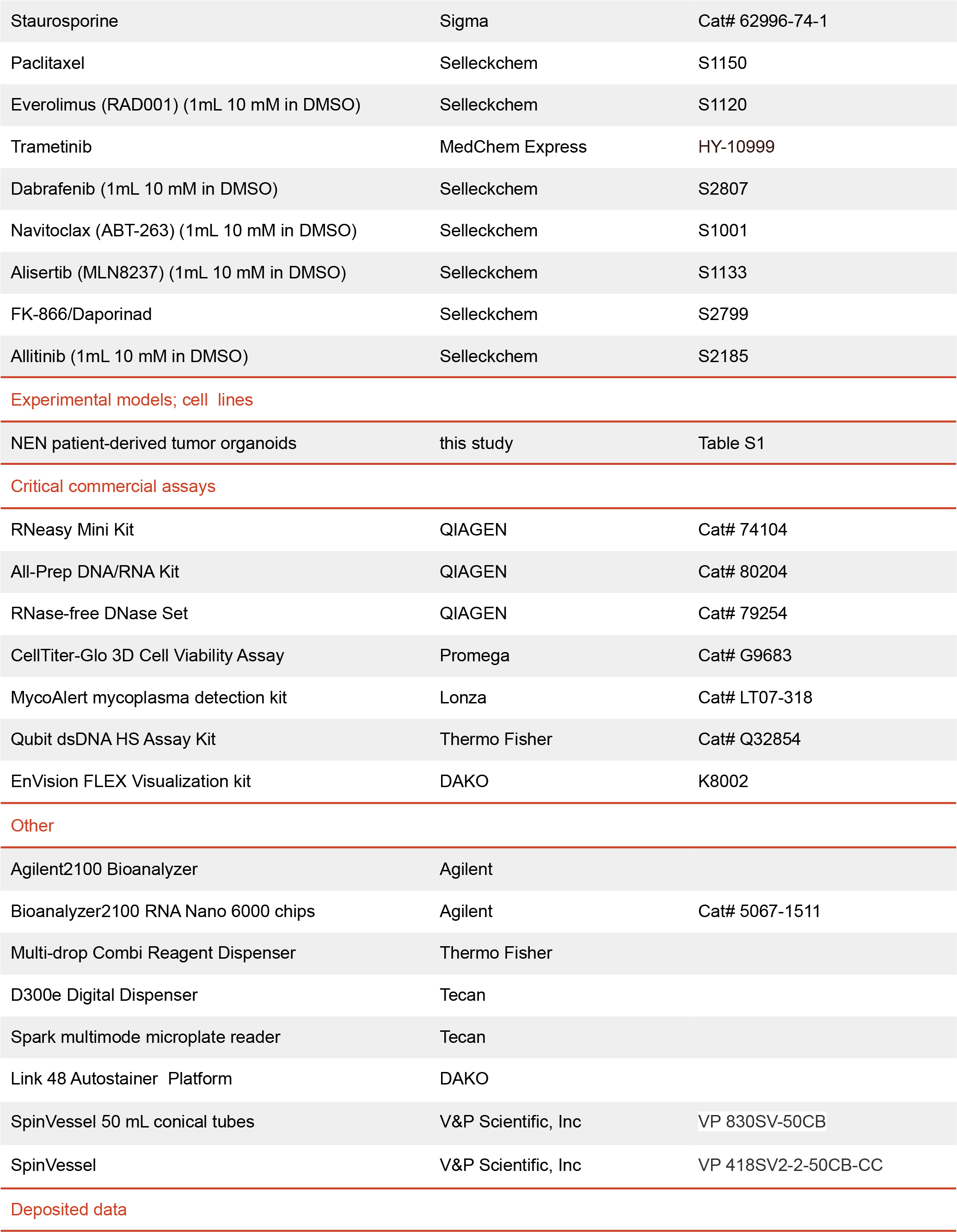
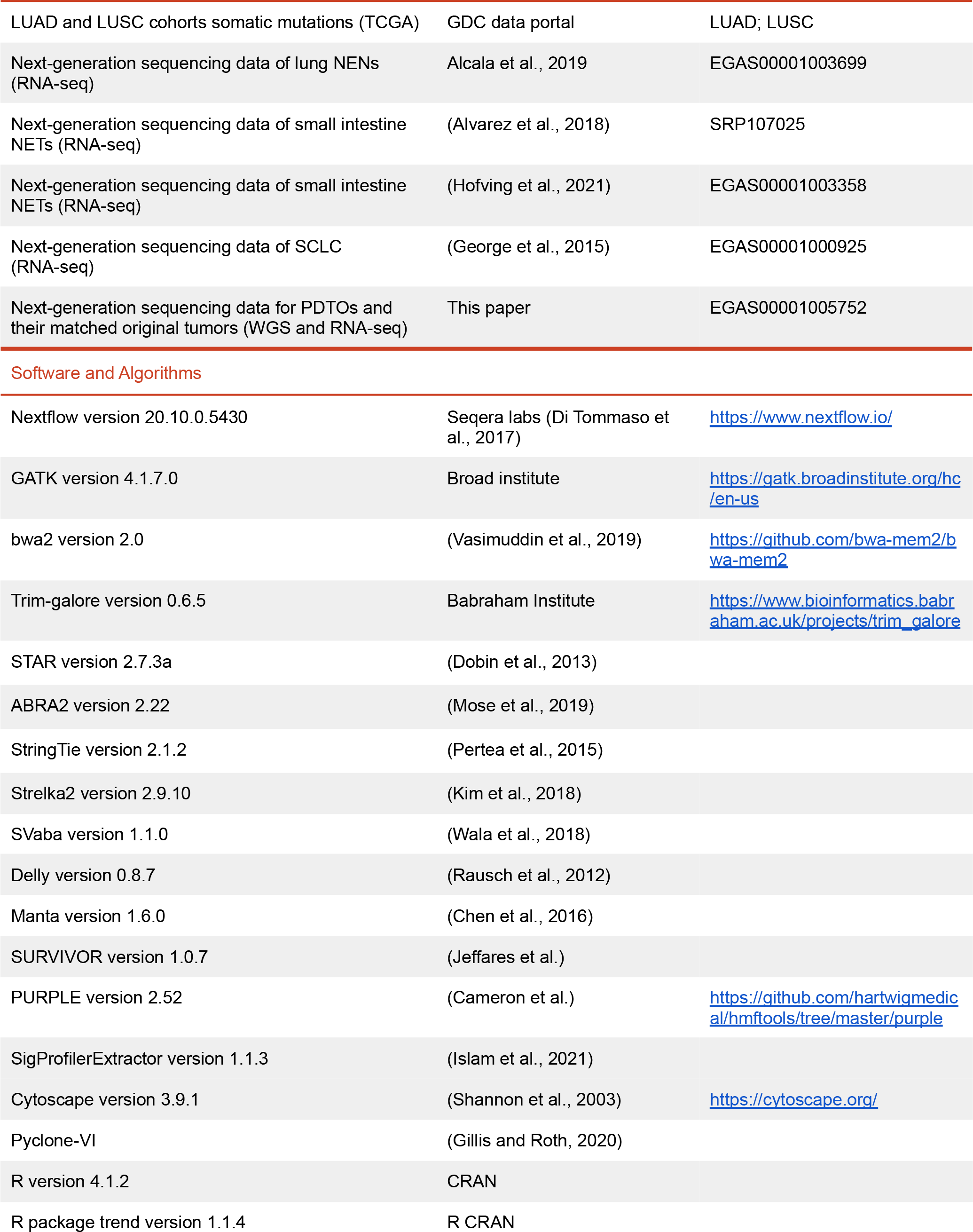
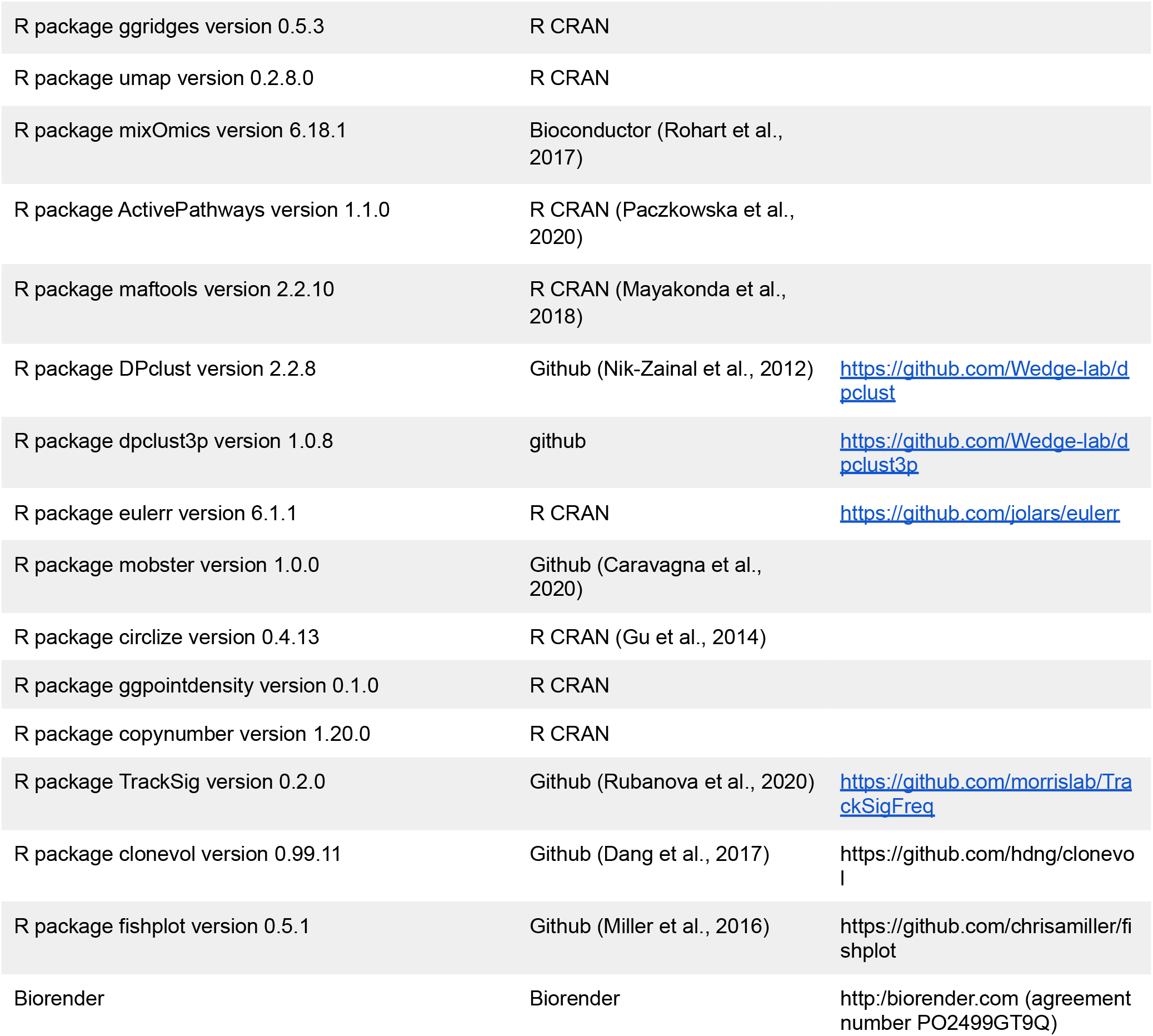

| REAGENT OR RESOURCE | SOURCE | IDENTIFIER |
| --- | --- | --- |
| Antibodies |  |  |
| Mouse monoclonal anti-Chromogranin A (LK2H10) | Thermo Scientific | MA5-13096 |
| Mouse monoclonal anti-human Ki67 antigen (MIB-1) | Agilent-Dako | GA62661-2 |
| Mouse monoclonal anti-human Synaptophysin | Dako | GA660 |
| Mouse monoclonal anti-human NCAM1/CD56 (clone 123C3) | Dako | IR628 |
| Rabbit monoclonal anti-EGFR (EP22) | Cell Marque | CMC41432010 |
| Chemicals, peptides, and recombinant proteins |  |  |
| DNase I | Sigma-Aldrich | Catalogue #DN25 |
| Collagenase | Sigma | C9407 |
| red blood cell lysis buffer | Roche | 11814389001 |
| Dispase II | Thermo Fisher | Cat# 17105041 |
| RNA<sup>Later</sup> Stabilization Solution | Thermo Fisher | AM7020 |
| Advanced DMEM/F12 | Thermo Fisher scientific | 12634-010 |
| DMEM + GlutaMAX | Thermo Fisher scientific | 31966-021 |
| B-27 Supplement | Thermo Fisher scientific | 17504044 |
| GlutaMAX | Thermo Fisher scientific | 35050061 |
| HEPES | Thermo Fisher scientific | 15630080 |
| Penicillin-Streptomycin | Thermo Fisher scientific | 15140122 |
| Noggin-Fc fusion protein | U-Protein Express | Cat# N002-100 ml |
| R-spondin3 conditioned medium | made in house |  |
| N-Acetyl-L-cysteine | Sigma-Aldrich | A1965 |
| Nicotinamide | Sigma-Aldrich | N0636 |
| Human EGF | Peprtech | AF-100-15 |
| A83-01 | Tocris | 2939 |
| Prostaglandin E2 | Tocris | 2296 |
| SB 202190 (p32 inhibitor) | Sigma-Aldrich | S7076 |
| Y-27632 dihydrochloride (ROCK inhibitor) | Abmole | M1817 |
| Primocin | Invivogen | ant-pm-2 |
| Cultrex Basement Membrane Extract (BME), Growth Factor Reduced, Type 2 | R&D Systems, Bio-Techne | 3533-001-02 |
| Formaldehyde Solution 4% | Sigma-Aldrich | 1.00496 |
| TrypLE Express | Thermo Fisher Scientific | 12605010 |
| CHIR (Chir99021) | Stemgent | Cat# 04-0004-10 |
| FGF2 | Peprtech | Cat# 100-18B |
| FGF7 | Peprtech | Cat# 100-19 |
| FGF10 | Peprtech | Cat# 100-26 |
| Staurosporine | Sigma | Cat# 62996-74-1 |
| Paclitaxel | Selleckchem | S1150 |
| Everolimus (RAD001) (1mL 10 mM in DMSO) | Selleckchem | S1120 |
| Trametinib | MedChem Express | HY-10999 |
| Dabrafenib (1mL 10 mM in DMSO) | Selleckchem | S2807 |
| Navitoclax (ABT-263) (1mL 10 mM in DMSO) | Selleckchem | S1001 |
| Alisertib (MLN8237) (1mL 10 mM in DMSO) | Selleckchem | S1133 |
| FK-866/Daporinad | Selleckchem | S2799 |
| Allitinib (1mL 10 mM in DMSO) | Selleckchem | S2185 |
| Experimental models; cell lines |  |  |
| NEN patient-derived tumor organoids | this study | Table S1 |
| Critical commercial assays |  |  |
| RNeasy Mini Kit | QIAGEN | Cat# 74104 |
| All-Prep DNA/RNA Kit | QIAGEN | Cat# 80204 |
| RNase-free DNase Set | QIAGEN | Cat# 79254 |
| CellTiter-Glo 3D Cell Viability Assay | Promega | Cat# G9683 |
| MycoAlert mycoplasma detection kit | Lonza | Cat# LT07-318 |
| Qubit dsDNA HS Assay Kit | Thermo Fisher | Cat# Q32854 |
| EnVision FLEX Visualization kit | DAKO | K8002 |
| Other |  |  |
| Agilent2100 Bioanalyzer | Agilent |  |
| Bioanalyzer2100 RNA Nano 6000 chips | Agilent | Cat# 5067-1511 |
| Multi-drop Combi Reagent Dispenser | Thermo Fisher |  |
| D300e Digital Dispenser | Tecan |  |
| Spark multimode microplate reader | Tecan |  |
| Link 48 Autostainer Platform | DAKO |  |
| SpinVessel 50 mL conical tubes | V&P Scientific, Inc | VP 830SV-50CB |
| SpinVessel | V&P Scientific, Inc | VP 418SV2-2-50CB-CC |
| Deposited data |  |  |
| LUAD and LUSC cohorts somatic mutations (TCGA) | GDC data portal | LUAD; LUSC |
| Next-generation sequencing data of lung NENs (RNA-seq) | Alcala et al., 2019 | EGAS00001003699 |
| Next-generation sequencing data of small intestine NETs (RNA-seq) | (Alvarez et al., 2018) | SRP107025 |
| Next-generation sequencing data of small intestine NETs (RNA-seq) | (Hofving et al., 2021) | EGAS00001003358 |
| Next-generation sequencing data of SCLC (RNA-seq) | (George et al., 2015) | EGAS00001000925 |
| Next-generation sequencing data for PDTOs and their matched original tumors (WGS and RNA-seq) | This paper | EGAS00001005752 |
| Software and Algorithms |  |  |
| Nextflow version 20.10.0.5430 | Seqera labs (Di Tommaso et al., 2017) | https://www.nextflow.io/ |
| GATK version 4.1.7.0 | Broad institute | https://gatk.broadinstitute.org/hc/en-us |
| bwa2 version 2.0 | (Vasimuddin et al., 2019) | https://github.com/bwa-mem2/bwa-mem2 |
| Trim-galore version 0.6.5 | Babraham Institute | https://www.bioinformatics.babraham.ac.uk/projects/trim_galore |
| STAR version 2.7.3a | (Dobin et al., 2013) |  |
| ABRA2 version 2.22 | (Mose et al., 2019) |  |
| StringTie version 2.1.2 | (Pertea et al., 2015) |  |
| Strelka2 version 2.9.10 | (Kim et al., 2018) |  |
| SVaba version 1.1.0 | (Wala et al., 2018) |  |
| Delly version 0.8.7 | (Rausch et al., 2012) |  |
| Manta version 1.6.0 | (Chen et al., 2016) |  |
| SURVIVOR version 1.0.7 | (Jeffares et al.) |  |
| PURPLE version 2.52 | (Cameron et al.) | https://github.com/hartwigmedical/hmftools/tree/master/purple |
| SigProfilerExtractor version 1.1.3 | (Islam et al., 2021) |  |
| Cytoscape version 3.9.1 | (Shannon et al., 2003) | https://cytoscape.org/ |
| Pyclone-VI | (Gillis and Roth, 2020) |  |
| R version 4.1.2 | CRAN |  |
| R package trend version 1.1.4 | R CRAN |  |
| R package ggridges version 0.5.3 | R CRAN |  |
| R package umap version 0.2.8.0 | R CRAN |  |
| R package mixOmics version 6.18.1 | Bioconductor (Rohart et al., 2017) |  |
| R package ActivePathways version 1.1.0 | R CRAN (Paczkowska et al., 2020) |  |
| R package maftools version 2.2.10 | R CRAN (Mayakonda et al., 2018) |  |
| R package DPclust version 2.2.8 | Github (Nik-Zainal et al., 2012) | https://github.com/Wedge-lab/dpclust |
| R package dpclust3p version 1.0.8 | github | https://github.com/Wedge-lab/dpclust3p |
| R package eulerr version 6.1.1 | R CRAN | https://github.com/jolars/eulerr |
| R package mobster version 1.0.0 | Github (Caravagna et al., 2020) |  |
| R package circlize version 0.4.13 | R CRAN (Gu et al., 2014) |  |
| R package ggpointdensity version 0.1.0 | R CRAN |  |
| R package copynumber version 1.20.0 | R CRAN |  |
| R package TrackSig version 0.2.0 | Github (Rubanova et al., 2020) | https://github.com/morrislab/TrackSigFreq |
| R package clonevol version 0.99.11 | Github (Dang et al., 2017) | https://github.com/hdng/clonevol |
| R package fishplot version 0.5.1 | Github (Miller et al., 2016) | https://github.com/chrisamiller/fishplot |
| Biorender | Biorender | http://biorender.com (agreement number PO2499GT9Q) |

### RESOURCE AVAILABILITY

#### Lead contact

Further information and requests for resources and reagents should be directed to and will be fulfilled by the Lead Contact, Hans Clevers or by Talya Dayton.

#### Data and code availability

The raw and processed next-generation sequencing data generated for this study will be deposited on the European Genome-phenome Archive (EGA). All bioinformatic processing pipelines are open-source and freely accessible via Github at https://github.com/IARCbioinfo/. Analysis scripts used to analyze the sequencing data and produce the related figures are available via Github at https://github.com/IARCbioinfo/MS_panNEN_organoids.

### METHOD DETAILS

#### Approval of studies involving human and patient-informed consent

The collection of patient data and tissue for the generation and distribution of normal and NEN organoids was performed according to the guidelines of the European Network of Research Ethics Committees (EUREC) following European, national and local law. The protocols were approved by the medical ethical committee (METC) corresponding to the respective hospitals where patients were treated: Verenigde Commissies Mensgebonden Onderzoek of the St. Antonius Hospital Nieuwegein, Z-12.55; UMC Utrecht, METC 12-093 HUB-Cancer; NKI Institutional Review Board (IRB), M18ORG/CFMPB582; Maastricht University Medical Center, METC 2019-1061, and 2019-1039. All patients participating in this study signed informed consent forms and could withdraw their consent at any time.

Organoid lines reported in this manuscript can be requested from the lead author, and/or from. Distribution of organoids to third parties will have to be authorized by the relevant ethical committee and a complete MTA will be required to ensure compliance with the Dutch ‘medical research involving human subjects’ act. Use of organoids is subjected to patient consent; upon consent withdrawal, distributed organoid lines and any derived material will have to be promptly disposed of.

#### NEN tissue processing

On arrival, NEN tissues were cut into 3–5 mm^3^ pieces. Two or three random pieces were placed in RNAlater solution and stored at −80 °C for DNA and RNA isolation, one random piece was fixed in formalin for histopathological analysis and immunohistochemistry, and the remainder were processed for organoid derivation. For organoid derivation: tissue was minced, collected with 10 mL DMEM containing antibiotics, and transferred to a 15 mL conical tube. To dissociate the minced tissue further, 200 μL of collagenase solution (20 mg ml−1) was added to the tissue/DMEM solution and the tube was placed on an orbital shaker at 37 °C for 25 minutes. After digestion, 50 μL of DNase I solution (10mg/mL) was added. The digested tissue suspension was sheared using 5 mL plastic pipettes, strained over a 100 μm filter. Large tissue pieces that remained after digestion were presumed to be necrotic or fibrotic and discarded. The filtered tissue suspension was centrifuged at 1,000 r.p.m. and the supernatant was removed. In case of a visible red pellet, erythrocytes were lysed in 50 to 300 μL of red blood cell lysis buffer (depending on pellet size) for 5 min at room temperature and then washed twice with 13 mL DMEM containing antibiotics. Finally, the pellet was resuspended in appropriate volume of BME for plating in 30 μL droplets on preheated suspension plates (Greiner).

#### Tumor organoid culture

NEN patient-derived tumor organoids were grown in 30 μL drops of BME in suspension culture plates, overlaid with growth medium. The growth medium consisted of Advanced DMEM/F12 supplemented with 1x GlutaMAX, 10 mM HEPES, penicillin-streptomycin, Primocin (InvivoGen, Cat# N001), 1% Noggin conditioned medium (U-Protein Express, Cat# N002), 20% of RSPO3 conditioned medium (made in-house), 1x B27 supplement (GIBCO, Cat# 175044), 1.25 mM n-Acetylcystein (Sigma, Cat# A9165), 3 μM CHIR (Stemgent, Cat# 04-0004-10), 1 μM Prostaglandin E2 (Tocris, Cat# 2296), 0.005 μg/mL FGF2 (Peprotech), 10 μM ROCK inhibitor (Abmole, Cat# Y27632), 500 nM A83-01 (Tocris, Cat# 2939), 3 μM p38 inhibitor SB202190 (Sigma, Cat# 7067). All lung NET organoids and some LCNEC organoids were grown in media additionally supplemented with 0.05 μg/ml EGF (Peprotech, Cat# AF-100-15). Media was changed once a week.

For splitting, a 2 minute incubation in TrypLE at 37°C was followed by mechanical shearing through a p10 tip attached a fire-polished plugged glass pipettes (Fisher Scientific, Cat# 11506973) was used to break organoids up into small clusters of cells. Following isolation, LCNEC lines were consistently passaged at an approximate splitting ratio of 1:12 every 14 days. Pulmonary NET lines show a high degree of variability in their growth rates and required passaging at a split ratio of 1:2 or 1:3 every 2 month for lines derived from grade 2 tumors and once every 3 months of lines derived from grade 1 tumors. SI NET PDTOs showed a similar growth rate as pulmonary NET PDTOs but could not be passaged more than 4 times. All organoid lines tested negative in the MycoAlert mycoplasma detection kit (Lonza, LT07-318).

#### Histological analyses

Tissue and organoids were fixed overnight in 4% paraformaldehyde and subsequently dehydrated, paraffin embedded, and sectioned. Standard H&E staining was performed and stained sections were blindly analyzed by a pathologist specialized in NENs. WHO criteria for NENs were applied. Immunohistochemistry was performed using antibodies against Chromogranin A (Thermo Scientific, clone: LK2H10) dilution 1:1000, Ki67 clone MIB-1 (DAKO, ‘ready to use’), Synaptophysin (DAKO, ‘ready to use’), CD56/NCAM1 (DAKO, ‘ready to use’), and EGFR (Cell Marque, EP22, 1:200). All stainings were performed using the DAKO Link 48 Autostainer Platform and the EnVision FLEX Visualization kit (DAKO, cat# K8002) according to standard diagnostic routine protocols and manufacturer instructions. Immunohistochemical stainings were evaluated by pathologists (L. Brosens, or L.M. Hillen and S. Lantuejoul), who were blinded for all clinical, histopathological, and sequencing data. Slides were scanned using a Pannoramic 1000 slide scanner (3DHISTECH Ltd) and images were acquired using the CaseViewer software (3DHISTECH Ltd).

#### Tissue Microarray analysis

EGFR expression in human pulmonary neuroendocrine tumors was analyzed in two tissue microarrays (TMA) from the Maastricht University Medical Center (MUMC+), jointly containing cores derived from 70 tumors. Each tumor on the TMAs was represented by 3 cores derived from both central and peripheral tumor regions. Each TMA core was independently scored for EGFR IHC intensity and membrane localization and given a score of either high, medium, low, or absent EGFR membrane expression, where at least 20% of tumor cells were positive. In the majority of positive cases 70 to 100% of tumor cells were positive. Subsequently, tumors were assigned an overall score. In cases where cores from the same tumor were given a different EGFR IHC score, the tumor was assigned the score consistent between at least 2 out of the 3 cores (see Table S5).

#### RNA and DNA isolation

RNA was isolated from NEN organoids using the RNeasy Mini Kit (QIAGEN, Cat# 74104) following the manufacturer’s instructions including DNaseI treatment (QIAGEN, Cat# 79254). RNA and DNA were isolated from the same sample of NEN organoids and/or tissue using the All Prep DNA/RNA Mini Kit (QIAGEN, Cat # 80204) following the manufacturer’s instructions.

#### Classification of NEN PDTOs as “high-purity” or “mixed”

Established NEN PDTO lines were classified as either “high-purity” or “mixed” according to available data derived from: morphological/histological analyses and molecular analyses. The term “high-purity” was applied to PDTO lines for which there was no discernible contamination in the organoids from non-tumor cells as identified by either histology or molecular analyses. The term “mixed” was applied to PDTOs for which histology and/or molecular analyses provided evidence for the presence of both tumor cells and non-tumor cells. The following criteria were defined as evidence of tumor cells in the culture: 1) presence of neuroendocrine marker-expressing cells identified by immunohistochemical staining (CHGA, SYP, or CD56/NCAM); 2) levels of neuroendocrine marker expression within the range observed for corresponding subtype in reference dataset (Figures 3 and S3; Table S2); 3) UMAP clustering of organoid sample together with corresponding parental tumor tissue (Figures 3 and S3); 4) identification of shared genetic driver or NEN-associated alterations in PDTO and parental tumors by WGS (when available); 5) identification of genetic driver or NEN-associated alterations in RNAseq reads (applied when WGS data was not available) (Figures 4 and S4; Table S4).

In all cases, a sample was defined as “pure” by criteria 1 when 60% or more of cells were NE-marker positive, and “mixed” when less than 60% of cells were NE-marker positive. For lines that were defined as “mixed” by criteria 1, evidence of tumor cell content by criterias 2 - 5 were used to determine whether the NE-marker+ cells were likely to be tumor cells (i.e. criteria 2 - 5 were used to classify lines for which criteria 1 showed less than 60% of cells were NE-marker+). When WGS data was available, tumor purity was estimated jointly with the copy number alterations using software PURPLE (see “copy number variant calling” paragraph below). One sample, LNET2Np7, derived from tumor adjacent normal tissue of LNET2 patient was assessed using this criteria. However, WGS analyses were inconclusive with regards to tumor cell content. Given the concurrent diagnosis of diffuse idiopathic pulmonary neuroendocrine cell hyperplasia (DIPNECH) in this patient and the high likelihood of DIPNECH cells in the “normal adjacent tissue,” this sample was removed from downstream analyses.

#### Nicotinamide assays

Single cells (4,000) were plated in 5 μL of BME in the wells of a 96 well plate and overlaid with media containing different concentrations of nicotinamide. Following expansion and organoid formation (one to four weeks depending on the line), ATP levels were measured using the CellTiter-Glo 3D Reagent (Promega, catalog no. G9681) according to the manufacturer’s instructions, and luminescence was measured using a Spark multimode microplate reader (Tecan). Results for each line were normalized to the value for that line in 10 mM nicotinamide, the standard concentration used in organoid culture (100%).

#### Quantitation of EGF dependency in lung NET organoids

Single cells (4,000) were plated in 5 μL of BME in the wells of a 96 well plate and overlaid with media containing different concentrations of EGF. Following expansion and organoid formation (one to ten weeks depending on the line), ATP levels were measured using the CellTiter-Glo 3D Reagent (Promega, catalog no. G9681) according to the manufacturer’s instructions, and luminescence was measured using a Spark multimode microplate reader (Tecan). Results for each line were normalized to the value for that line in media lacking EGF (100%). Values shown in the graph were then normalized across rows by dividing by the highest viability value.

#### Drug sensitivity tests

Prior to the start of the drug screen, TrypLE was used to disrupt organoids into single cells and small clusters of cells that were then plated in 30 μL drops BME in the wells of a 6 well plate. Organoids were then grown in the appropriate media as for regular expansion. Seven to ten days later, organoids were collected from the BME by the addition of 1 mg/mL dispase II (Sigma-Aldrich, catalog no. D4693) to the medium of the organoids. Organoids were incubated for 90 minutes at 37°C to digest the BME. Following collection of organoids and washing with DMEM to remove dispase, organoids were filtered using a 70-mm nylon cell strainer (BD Falcon), counted, and resuspended in 5% BME/ growth medium (12,500 organoids/mL) and transferred to SpinVessel 50 mL conical bottom tubes. The SpinVessel tubes containing the organoid solution were placed on a SpinVessel machine and rotated at speed setting 25 so as to keep organoids in a homogenous solution. Finally, 40 μL volume of organoid solution was dispensed (Multidrop Combi Reagent Dispenser, Thermo Scientific, catalog no. 5840300) in 384-well plates (Corning, catalog no. 4588).

Drugs were added 3 days after plating using the Tecan D300e Digital Dispenser (Tecan). The time in between plating and addition of drugs was maintained to allow organoids to recover after plating. All drugs in the assays were dissolved in DMSO and all wells were normalized for DMSO percentage, which never exceeded 1%. Drug exposure was performed in triplicate for each concentration shown and drug assays were repeated at least once for all drugs. When available drugs were purchased as 1 mL of a 10 mM solution in DMSO (See reagents table). For FK866 5 mM solution was made in DMSO and further diluted to reach the assay concentrations. 10 mM solutions in DMSO were peprared of staurosporin, trametinib, and paclitaxel.

Seven days after adding the drugs, ATP levels were measured using the CellTiter-Glo 3-D Reagent (Promega, catalog no. G9681) according to the manufacturer’s instructions, and luminescence was measured using a Spark multimode microplate reader (Tecan). Results were normalized to vehicle (100%) and baseline control (Staurosporin 1 μmol/L; 0%). For each line, when viability did not go above 70% or below 30%, an additional screen was performed for that particular drug with an adjusted dose of this drug for this organoid line. Screen quality was determined by checking *Z* factor scores for each plate following this formula:

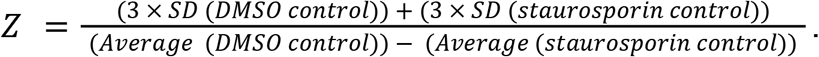

The average *Z* factor score for all assays included in the manuscript was 0.7 (ranging from 0.40 to 0.81), which is consistent with an experimentally robust assay.

#### Statistical analyses

Temporal trends in passage times were tested using Mann-Kendall trend tests as implemented in the R package trend, for each sample individually. Resulting *p*-values were adjusted for multiple testing using the Benjamini-Hochberg procedure (Benjamini and Hochberg, 1995).

#### Whole-genome sequencing

*Lab work*. DNA was isolated from tumor tissue and organoid samples using the All Prep DNA/RNA Mini Kit (QIAGEN, Cat # 80204) following the manufacturer’s instructions. For a blood germline reference, isolation of DNA from blood samples donated by corresponding patients was performed by USEQ (Utrecht Sequencing Facility) using the QIAGEN QIASymphony SP. Quality and quantification of DNA samples were checked with Qubit (DNA BR). DNA integrity and RNA contamination was assessed by using Tapestation DNA screens (Genomic screen) and Nanodrop (260/280 ratio). Per sample, 500–1,000 ng of DNA was used for DNA library preparation by USEQ using the TruSeq DNA Nano kit. Whole-genome sequencing was performed on NovaSeq 6000 to an average coverage of 30x for germline reference samples, and 60x to 90x for tumor tissue and organoid samples (see Table S4).

Early passage was defined as passages 1 to 3, intermediate passage as 4 to 6, and late passage as passages 7 and beyond.

*Processing*. Raw reads were processed using our in-house whole-genome sequencing processing workflow https://github.com/IARCbioinfo/alignment-nf v 1.2, which uses software bwa to align reads to reference genome GRCh38 with decoy genome and alt contigs, and GATK to perform base quality score recalibration. We performed quality controls using fastQC for raw reads and qualimap for aligned reads (using workflow https://github.com/IARCbioinfo/qualimap-nf v. 1.1), and confirmed that files of a same experiment came from the same individual using NGSCheckMate (using workflow https://github.com/IARCbioinfo/NGSCheckMate-nf v. 1.1a).

#### RNA-sequencing

##### Lab work

We performed paired-end bulk RNA sequencing (RNA-seq) across 20 patients, including 1 patient with both primary tumor and metastasis, for a total of 18 parental tumor samples and 21 NEN PDTO lines that had been in culture for 6 to 67 weeks (Table S2). For 5 PDTO lines we captured expression patterns from 2 separate time points in culture. RNA was isolated as described above. RNA integrity was measured using the Agilent RNA 6000 Nano kit with the Agilent 2100 Bioanalyzer and RNA concentrations were determined using the Qubit RNA HS Assay Kit. RIN values of organoid RNA samples were typically 9-10 and only samples with RIN >8 were used for library preparation. RIN values of tissue RNA samples were typically 8-10, with 5 samples showing an RIN between 6.3 and 7.4, and only samples with RIN >6 were used for library preparation. RNA libraries were prepared using TruSeq Stranded mRNA polyA kit (Illumina) and sequenced on either an Illumina Nextseq 2000 (paired-end, 2 x 150 bp) or an Illumina Novaseq 6000 (paired-end, 2 x 150 bp). Library preparation and sequencing was performed by USEQ (Utrecht Sequencing Facility).

##### Processing

Processing of 47 RNA-seq from the experiments and public data for 210 RNA-seq from LNEN (Alcala et al., 2019), 30 LNETs (Laddha et al., 2019), and 88 RNA-seq from SINET (Alvarez et al., 2018; Hofving et al., 2021) were all done using our RNA-seq pre-processing workflow (https://github.com/IARCbioinfo/RNAseq-nf v. 2.4) to ensure mitigation of potential batch effects between cohorts due to differences in processing (software, version, or operating system); as described in Gabriel et al. (2020), this workflow uses trimgalore to trim reads for adapter sequences and STAR to map reads to reference genome GRCh38. Base quality scores were then recalibrated to improve subsequent variant calling using GATK (https://github.com/IARCbioinfo/BQSR-nf v. 1.1), and alignments were realigned locally using ABRA2 to improve their quality, in particular at splicing junctions (https://github.com/IARCbioinfo/abra-nf v. 3.0).

##### Expression Quantification

Gene expression quantification was performed with Stringtie (Pertea et al., 2015), in one-pass mode, using our workflow https://github.com/IARCbioinfo/RNAseq-transcript-nf v 2.2 with the gencode v33 comprehensive gene annotation as reference, providing expression in raw read count and TPM formats. We also processed the SINET RNA-seq datasets EGAS00001003358 and SRP107025 (Alvarez et al., 2018; Hofving et al., 2021) with the same worflows, which is also the version that was used to process the LNEN data by Gabriel and colleagues (Gabriel et al., 2020), allowing integration of the different datasets with minimal batch effects.

##### Unsupervised analyses

Analyses were performed separately on (i) lung and pancreatic NENs, and (ii) on small intestine NETs. Raw counts from all samples were normalized using the variance stabilization transform (R package DESeq2) to provide approximately normally distributed values, separately for (i) and (ii) and using the “blind” mode so each tumor type is processed in an independent and completely unsupervised manner (samples can have their own variance/mean relationship) but biologically meaningful differences between SINETs and other NENs are not removed. UMAP was then performed separately on (i) and (ii), using features known to be informative about molecular groups, and setting the number of neighbors to the maximal value (number of samples) so that both short- and long-distance relationships between samples are preserved, as in our recent integrative study (Gabriel et al., 2020). The subset of genes for the lung and pancreatic NENs (i) were extracted from a published list of “core” genes that are differentially expressed between all pairs of lung NEN molecular groups (Alcala et al., 2019), excluding genes that were discarded between gencode v19 and v33 or that changed ENSEMBL ID between the releases (54/1459 genes). Cluster annotations were extracted from Gabriel et al. (2020) (https://github.com/IARCbioinfo/DRMetrics/data/Attributes.txt.zip), assuming as shown in the article that clusters A1, A2, and B from Alcala et al. (2019) respectively correspond to clusters LC1, LC3, and LC2 from Simbolo et al. (2019), and grouping together under the term LCNEC (resp. SCLC) LCNEC and LCNEC-like SCLC (resp. SCLC and SCLC-like LCNEC). The subset of genes for the SINET samples (ii) included all 520 genes from Alvarez et al. (2018) except *NME1-NME2*, which is a readthrough transcription between neighboring genes *NME1* and *NME2* that was not quantified by StringTie.

##### Supervised analyses

A Partial Least Squares (PLS) analysis was performed on matched parental tumors (“parent” group) and PDTOs (“organoid” group) in order to find genes whose expressions are altered by the PDTO formation process, using the pls function from R package mixOmics in “regression” mode and selecting the first 10 components of the expression matrix (named *X* in the PLS framework, while the group matrix is *Y*), after running the variance stabilization transform on the entire dataset simultaneously using a design comparing the “parent” and “PDTO” groups, so differences between the groups is not removed during the transformation. PLS components separating parents and PDTOs were identified using ANOVA *F*-tests with each of the ten components as a function of the groups. We found only two components associated with the groups (F-tests, component 1, *p*<2.2✕10^−16^, component 2, *p*=0.01293, other components *p*>0.35).

To understand which genes were responsible for the separation between parents and PDTOs, we computed the Pearson correlation between the expression of each gene and each of the two PLS components associated with the groups and performed integrated gene set analysis on the correlation *p*-values using the ActivePathways R package with GO terms as gene sets. ActivePathways allows combining *p*-values from multiple sources (here components 1 and 2) to find which pathways are associated with each source separately and in combination. We found 108 significantly enriched pathways (see results in Fig. S3G and Table S2). Pathways were then aggregated into super-pathways using the EnrichmentMap module of cytoscape, which forms a network of pathways based on their similarity in terms of shared genes, using the default cutoff of 0.375 similarity to connect two pathways. Results are presented Fig. S3H, and show a vast majority (89/108) of pathways belonging to immune-related pathways and pathways sharing genes with these immune-related pathways. We also found two additional super pathways–one comprising four blood-vessel-related pathways and one comprising five synaptic-related pathways–a small group of 3 pathways and 6 additional isolated pathways.

#### Somatic alteration calling

##### Small variants calling

Single nucleotide variants were called from WGS data using software Mutect2 from GATK4 (using our workflow https://github.com/IARCbioinfo/mutect-nf v. 2.2b), and indels were called using the intersection of Mutect2 and strelka2 calls (using our workflow https://github.com/IARCbioinfo/strelka2-nf v. 1.2a) in order to filter out false positive calls, more numerous in indels than in SNVs. Annotations were performed with ANNOVAR (using our workflow https://github.com/IARCbioinfo/table_annovar-nf v. 1.1.1). To ensure comparisons with previous studies on NETs (Alcala et al., 2019; Fernandez-Cuesta et al., 2014) and common cancers (Hoadley et al., 2018), that mostly relied on whole-exome sequencing, tumor mutational burdens were computed using the exonic ranges from the SureSelect Human All Exon v7 panel from Agilent (bed file downloaded from the manufacturer’s website, approximately covering 36Mb), and focusing on non silent mutations (nonsnononymous SNVs, nonsense mutations, nonstop mutations, and indels).

Single nucleotide variants were called from RNA-seq data in the samples without WGS data using software Mutect2 from GATK4 as for WGS, but in tumor-only mode (also using our workflow https://github.com/IARCbioinfo/mutect-nf RNAseq branch) and annotated using ANNOVAR as described above. In addition, calls were further stringently filtered to reduce false somatic calls: we excluded variants reported with a population frequency >1% in any of the populations from database GNOMAD (ANNOVAR column “non_cancer_AF_popmax”), in ExAC excluding TCGA samples (ANNOVAR column ExAC_nontcga_ALL), or in the 1000 genomes project (ANNOVAR column 1000g2015aug_all), we excluded nonsynonymous variants with a REVEL pathogenicity score <0.7, and we excluded variants never reported in the COSMIC v92 catalog of coding cancer variants. Whenever a variant was found in either parental tumor or a PDTO, the corresponding position in the matched PDTO or parental tumor was classified as “no_coverage” whenever the read depth was 9 or less, below recommended depth for variant detection (Koboldt, 2020).

##### Structural Variant Calling

Structural variants (SVs) were called from WGS data using a two-step ensemble method combining results from 3 software: SVaba (Wala et al., 2018), Delly (Rausch et al., 2012), and Manta (Chen et al., 2016). In the discovery step, following Mangiante et al. (2021), we independently called SVs in each of the 23 tumor samples with WGS data using the SURVIVOR consensus calling (Jeffares et al.), merging SVs within a distance of 1kb and requiring either agreement between 2 SV callers or a strong support by a single caller (at least 15 reads), as implemented in our workflow (https://github.com/IARCbioinfo/sv_somatic_cns, v. 1.0). This step led to a list of high-quality SVs. In the subsequent recovery step, for each experiment and each SV that was not detected in all samples from this experiment, we checked whether one of the SV callers found reads supporting this SV with breakpoints within 1kb of those detected (SVs marked as “recovered” in Table S4), in unfiltered calls from SVaba and Manta, and initial calls from DELLY (no unfiltered calls are reported by the software). SVs were annotated based on their type (inversion, translocation, duplication, deletion), the position of their breakpoints (intergenic, within exons or introns) and their strands as described in Mangiante et al. (2021), using the gencode v33 annotation.

##### Copy Number Variant calling

Copy number variants (CNVs) were called from WGS data using software PURPLE using our custom workflow https://github.com/IARCbioinfo/purple-nf (branch dev_multi-sample). This workflow relies on the recommended PURPLE workflow using AMBER for B-allele frequency (BAF) estimation and COBALT for read depth ratio (RDR) estimation (https://github.com/hartwigmedical/hmftools/tree/master/purple), but relying on multi-sample segmentation of using the multipcf function from R package copynumber (Nilsen et al., 2012) instead of the simple pcf function used in the recommended workflow; this allows to infer more consistent breakpoints across samples from the same experiment. In addition, we used the option to include the list of SNVs called by Mutect2 (see above) to improve the CNV estimation by using the variant allelic fractions.

##### Analyses

We identified as damaging mutations the nonsynonymous mutations predicted as damaging by REVEL (score greater than or equal to 0.5) which combines multiple other damage predictions to maximize the evidence of pathogenicity (Ioannidis et al., 2016), along with stop gains, stop losses, indels, and splicing mutations. We retrieved driver gene lists from a literature search of lung and SI neuroendocrine neoplasms (Banck et al., 2013; Derks et al., 2018; Fernandez-Cuesta et al., 2014; George et al., 2018; Miyoshi et al., 2017; Pelosi et al., 2018; Rekhtman et al., 2016; Sei et al., 2015; Simbolo et al., 2017, 2018; Walter et al., 2018) (see Table S4 for the complete list). The waterfall plots (Figure 4B-C) were produced using R package maftools (Mayakonda et al., 2018). Circos plots were produced using R package circlize (Gu et al., 2014).

#### Mutational signatures

##### Small variants

Signatures were extracted from the VCFs using SigProfilerExtractor, comparing decompositions with 2 to 10 de novo signatures and using the number of signatures maximizing stability while minimizing cosine differences in signature reconstruction. Temporal signature activities were reconstructed using R package TrackSig separately on public (present in both parent and PDTO) and private alterations (present only in one sample). All signatures are reported in Fig. S5 and Table S4, and signatures known to be due to sequencing artifacts were removed and signature contributions normalized to sum to 100% for Fig. 5.

#### Evolutionary analyses

##### Mutation clustering and clonality

Somatic alterations were clustered and classified as clonal or subclonal using deconvolution of VAF distributions with R package DPclust (Nik-Zainal et al., 2012). Input files were prepared using R package dpclust3p, focusing on alterations with good coverage (above the target depth, 30X or 60X depending on the samples; see Table S4) and in clonal CNVs regions with consistent CN estimates in at least 2 samples from the same line for more accurate reconstructions. We used 5000 iterations after a burn-in phase of 1000 iterations. DPclust computed cancer cell fractions (CCFs) based on VAF and CN states and provided a likelihood that each alteration belongs to each cluster. In order to obtain high-confidence clonal reconstruction, we assigned alterations to a given cluster only if their likelihood to belong to this cluster was greater than or equal to 95%; other alterations were classified as “Uncertain” clustering. Clusters with less than 2.5% of assigned alterations were excluded. Clusters with an estimated location above a CCF of 0.95 in all samples were considered clonal. We summed up the likelihoods of belonging to each subclonal cluster, and considered alterations with a likelihood sum greater than or equal to 95% as subclonal. Venn-euler diagrams of shared clonal and subclonal alterations in Fig. 5(A) were computed using R package eulerr.

Clonality of driver small variant alterations not included in the DPclust model fit was subsequently assessed by finding the closest cluster in terms of CCF considering only alterations with the same CN as the focal driver variant. Clonality of CNVs was assessed using PURPLE estimated allelic fractions, with CNVs with both minor and major allele copy number estimates close to an integer value (with a threshold of 0.2) considered clonal, and the rest as subclonal. Clonality of structural variants was assessed by averaging the variant allelic fraction reported by each of the three callers and comparing with the expected clonal allelic fraction given the tumor purity. Clonality of the chromothripsis event in LNET10 was assessed using the clonality of the involved CNV segments (in particular in chr1 and chr9) and the allelic fractions of structural variants estimated from the proportion of supporting reads in the structural variants clustered in the involved regions (chr1, chr4, chr9, and chr16).

##### Phylogenies and fishplots

We used clonevol (Dang et al., 2017) to reconstruct tumor phylogenies between clusters. In order to account for the uncertainty in estimating the centroid of mutation clusters from CCF distributions, for each cluster we randomly sampled centroid positions from normal distributions centered on the estimated value and with a standard deviation equal to the observed standard deviation of CCF values for this cluster. We then ran clonevol independently on each randomly sampled centroid, and selected 20 resulting possible phylogenies; we selected the replicate with the CCF centroids closest to the point estimate for visualization. We plotted the results using R package fishplot (Miller et al., 2016). Because no clonal cluster and alterations were identified for LNET2, we used a polyclonal model; for all other samples, a clonal cluster was found and we used a monoclonal model. Note that alterations absent from a sample but that cluster with alterations at a non-zero frequency in this sample are thus inferred as part of a subclone detected in the focal sample; such mutations usually have a lower coverage in the focal sample, explaining the detection failure. In particular, although absent from sample LCNEC4 organoids, the *APC* and *PTPRZ1* mutations reported in Figure 4B were detected at a very low allelic fraction (6 and 7%, respectively) and were assigned to cluster 2 from Figure S5A, which is at low but non-zero frequency in the PDTOs (Figure 5B), and indeed coverage at their positions in the PDTOs were lower than in the original tumor (59 and 24 *vs* 89 for *APC*, and 96 and 24 *vs* 109 for *PTPRZ1*), making failure of detecting them if present in this cluster likely.

Similarly, the *ERCC6L* mutation in LNET2 and the *SMARCA4*, *ATR*, and *NTRK3* mutations were all assigned to low frequencies clones even though undetected in the corresponding sample.

##### Diversity summary statistics

Levels of intra-tumor genetic diversity were computed using the effective number of alleles *Δ* (Jost, 2008), a measure widely used in ecology and conservation genetics to monitor the level of genetic diversity in species of conservation interest (Jost et al., 2018). In the case of biallelic markers such as somatic small variants, *Δ* is a genetic diversity metric which ranges from 1 (no diversity) to 2 (maximal diversity) and captures how many alleles are effectively segregating in the population at a given polymorphic site; for example, given a polymorphic site with two alleles, A at a high frequency 0.999 and B at a low frequency 0.001, although two alleles are present, because allele B is only present in a fraction of individuals, the *effective* number of alleles is 1.002≃1. In order to obtain a quantity analogous to the tumor mutational burden, we rather focus on *Δ*-1, the effective number of *alternative* alleles (i.e., not taking into account the reference allele), and compute its sum across all polymorphic sites:

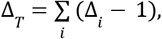

where 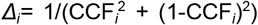 and CCF_*i*_ is the cancer cell fraction of the alternative allele at site *i*. We name this metric “effective number of alterations”, and further divide it by the size of the genomic ranges (in Mb) from which the alternative alleles were called. The resulting quantity ranges from 0, when there are no subclonal alterations or when all subclonal alterations have infinitesimal values, to the subclonal TMB when all clonal alterations are present in exactly half of the tumor cells (CCF=0.5, the situation maximizing diversity). All diversity statistics were computed using Nei and Chesser’s estimators (1983).

##### Mode of evolution

Neutral evolution was detected from the distribution of subclonal alterations using R package MOBSTER (Caravagna et al., 2020). MOBSTER uses a mixture model to identify a “neutral tail” in the distribution of CCF indicating the presence of a neutrally evolving subclone. For each sample, we filtered out alterations with a CCF below 5% and independently fitted models with and without neutral tails and with up to 4 additional subclones, and chose the best model using the reduced ICL (reICL) statistic.

---

**Figure S1.**
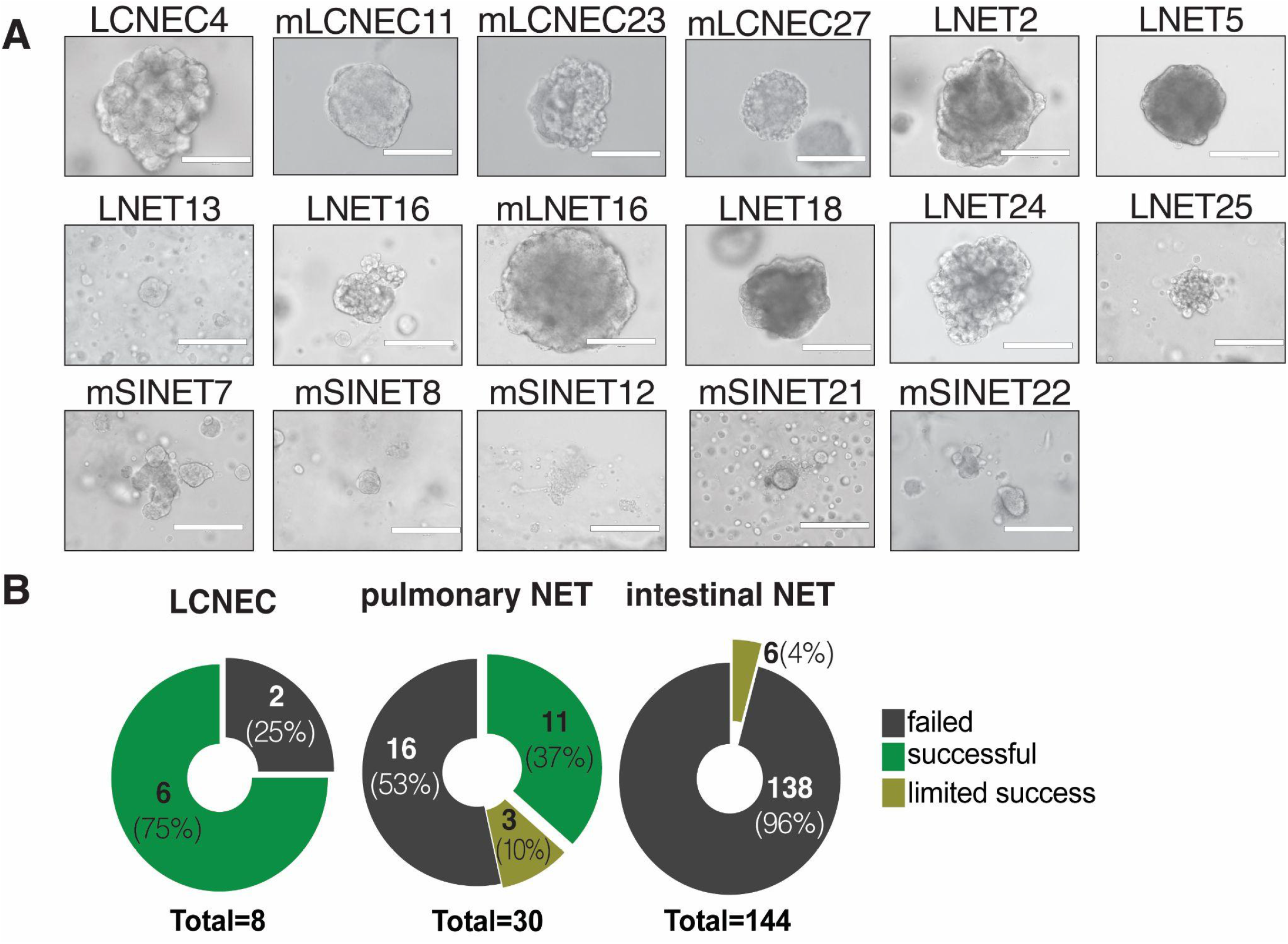
Establishment of a NEN PDTO biobank. (A) Representative bright-field images of NEN PDTOs. Scale bar: 200 μm except for mSINET12 where scale bar is 400 μm. (B) Success rate to establish PDTOs from isolated LCNEC, pulmonary NET, and intestinal NET tissue. Limited success indicates lines that were passaged long enough to generate molecular data but that subsequently stopped growing. Not shown: tissue from pancreatic NETs was also collected for this study but organoid generation was unsuccessful in all cases.

**Figure S2.**
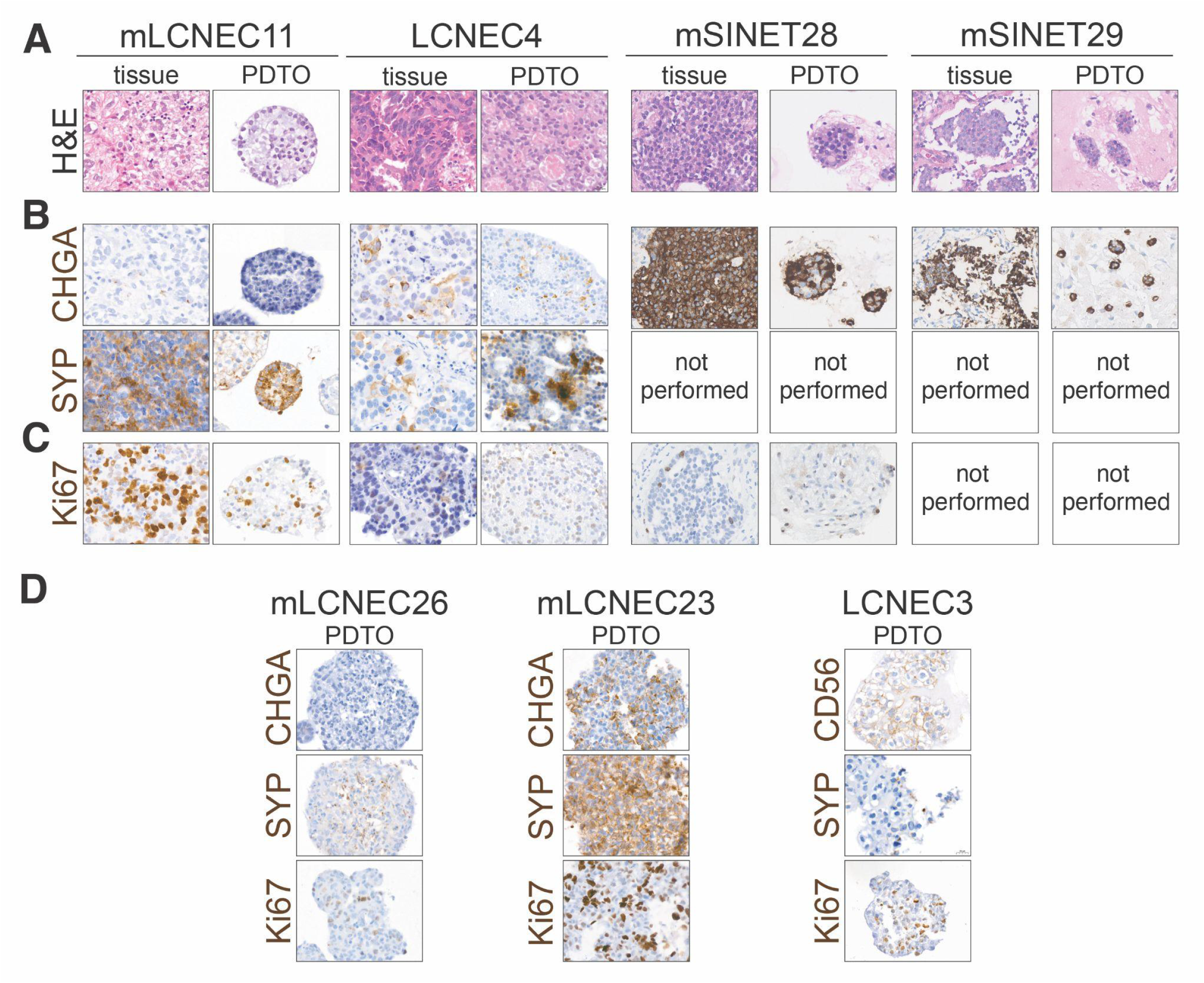
NET and LCNEC PDTOs retain histologic features of parental tumor subtypes. Representative images of (A) hematoxylin and eosin (H&E) staining and immunohistochemical staining for (B) the neuroendocrine marker, Chromogranin A (CHGA) and (C) the proliferation marker Ki67 of NEN PDTOs and their corresponding parental tumor tissue. Scale bar: 20 μm. mSINET: metastasis of small intestine NET; mLCNEC: metastasis of LCNEC. (D) Representative images of immunohistochemical staining for the neuroendocrine markers, Chromogranin A (CHGA) or CD56, and Synaptophysin (SYP), and the proliferation marker Ki67 of LCNEC PDTOs for which parental tissue was not available.

**Figure S3.**
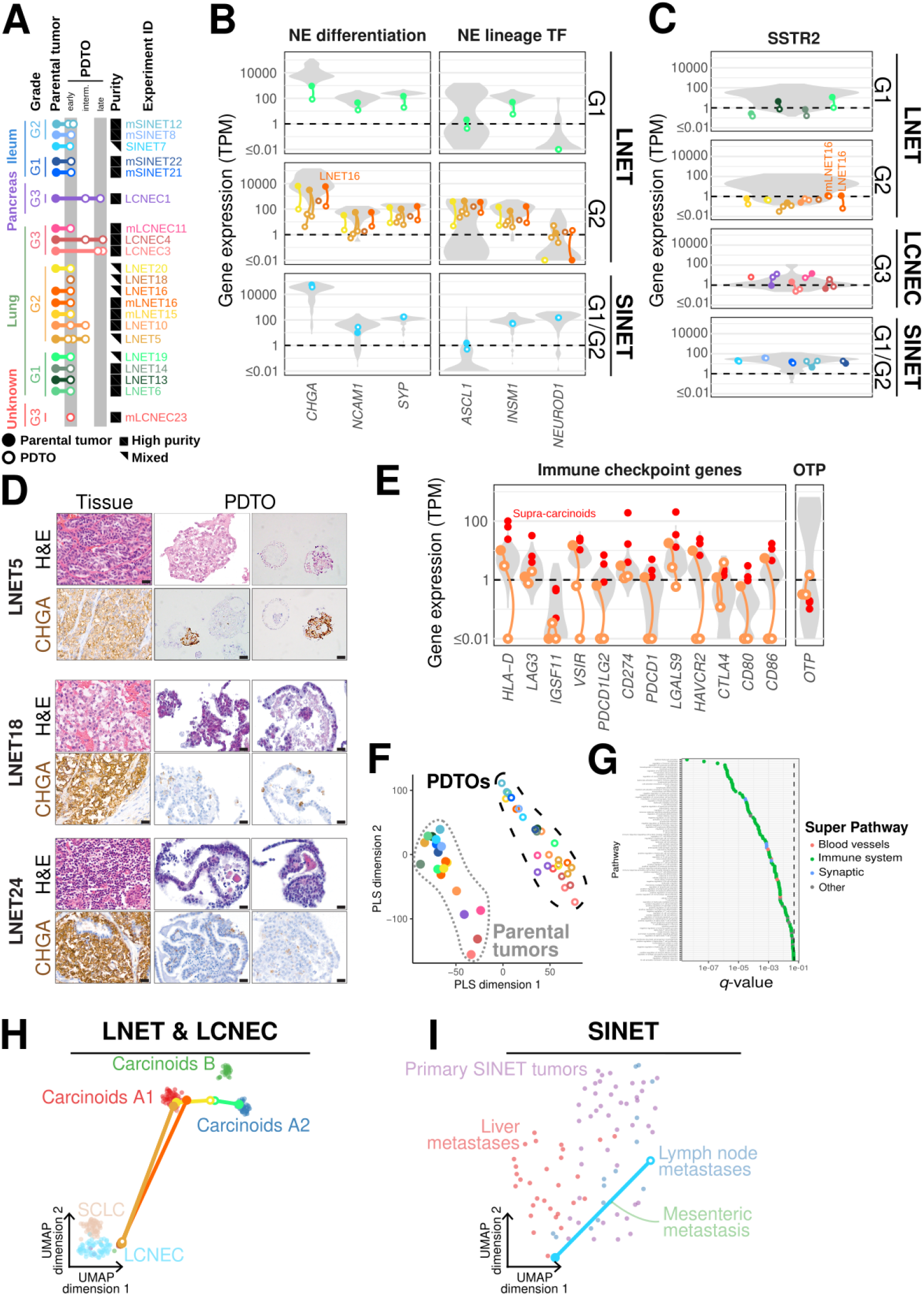
Expression analyses. (A) Extended panel 3A: outline of samples submitted to RNAseq (B) mRNA expression of neuroendocrine markers in mixed PDTOs and parental tumors. (C) SSTR2 mRNA expression in all samples. (D) Protein expression (IHC) for CHGA in low purity PDTOs (LNET5, LNET18, LNET24). Scale bar: 20 μm (E) mRNA expression of immune checkpoint genes and *OTP* in all samples. (F) PLS of PDTOs and tumors. (G) Gene set enrichment analysis of PLS components separating parental tumors and PDTOs. (H) UMAP of mixed LNET & LCNEC PDTOs. (I) UMAP of the mixed SINET PDTO. See **Table S2**.

**Figure S4.**
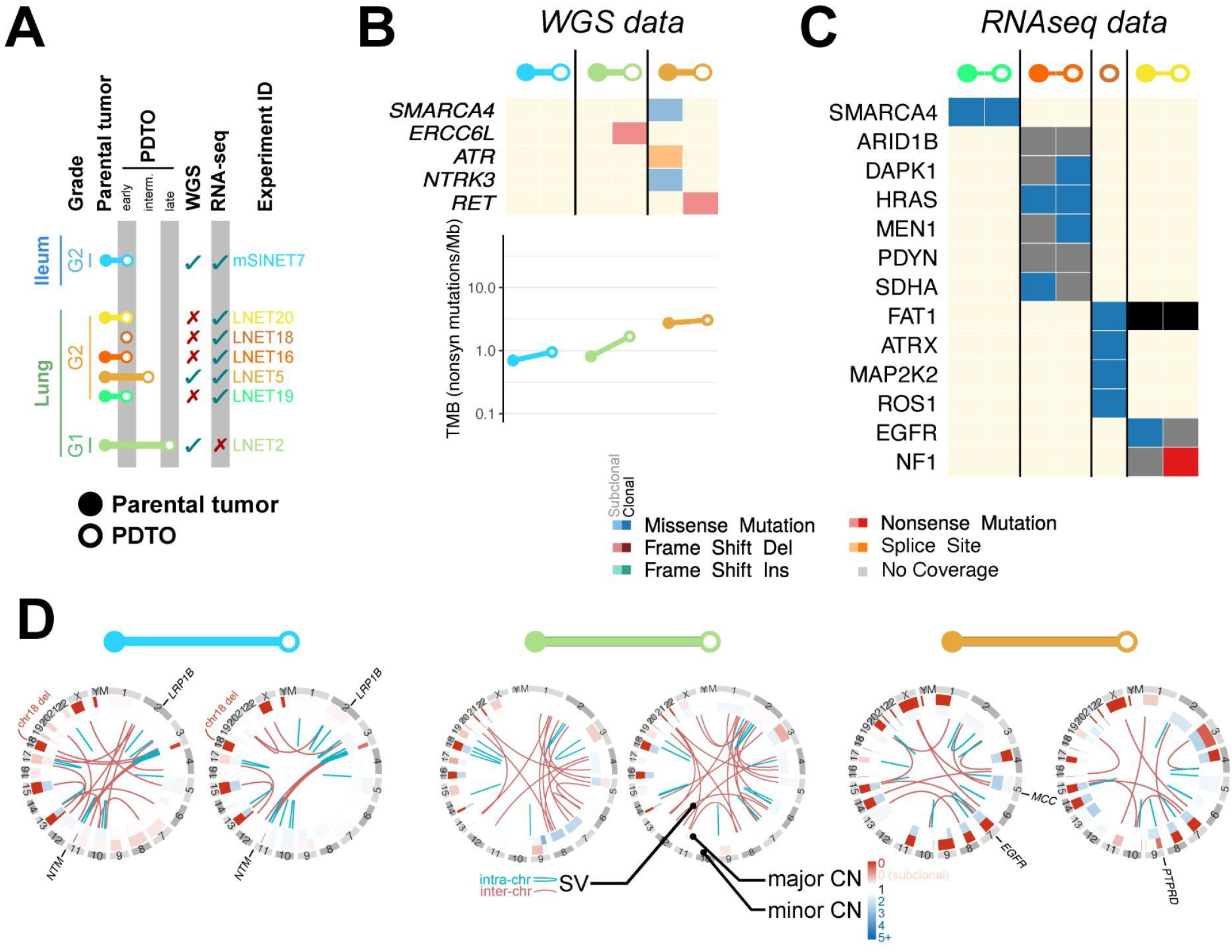
Genomic profiles of mixed PDTOs. All panels correspond to that of **Figure 4**, but focusing on mixed PDTOs SINET7, LNET2, and LNET5. In (C), note that the *EGFR* mutation in LNET20 is a non-recurrent mutation (T227C, COSMIC ID COSV51767338) only reported once in an acute lymphoblastic T cell leukemia tumor (Kalender Atak et al., 2012); in (D), the EGFR SV in LNET5 is an inversion of the 54,719,987–55,202,337 region (starting before exon 1 and ending in the intronic region between exons 26 and 27, out of the 28 exons of the main transcript ENST00000275493.7), with limited spanning and split reads support (13/126 and 8/123, respectively) suggesting a low-frequency subclonal alteration. See **Table S4**.

**Figure S5.**
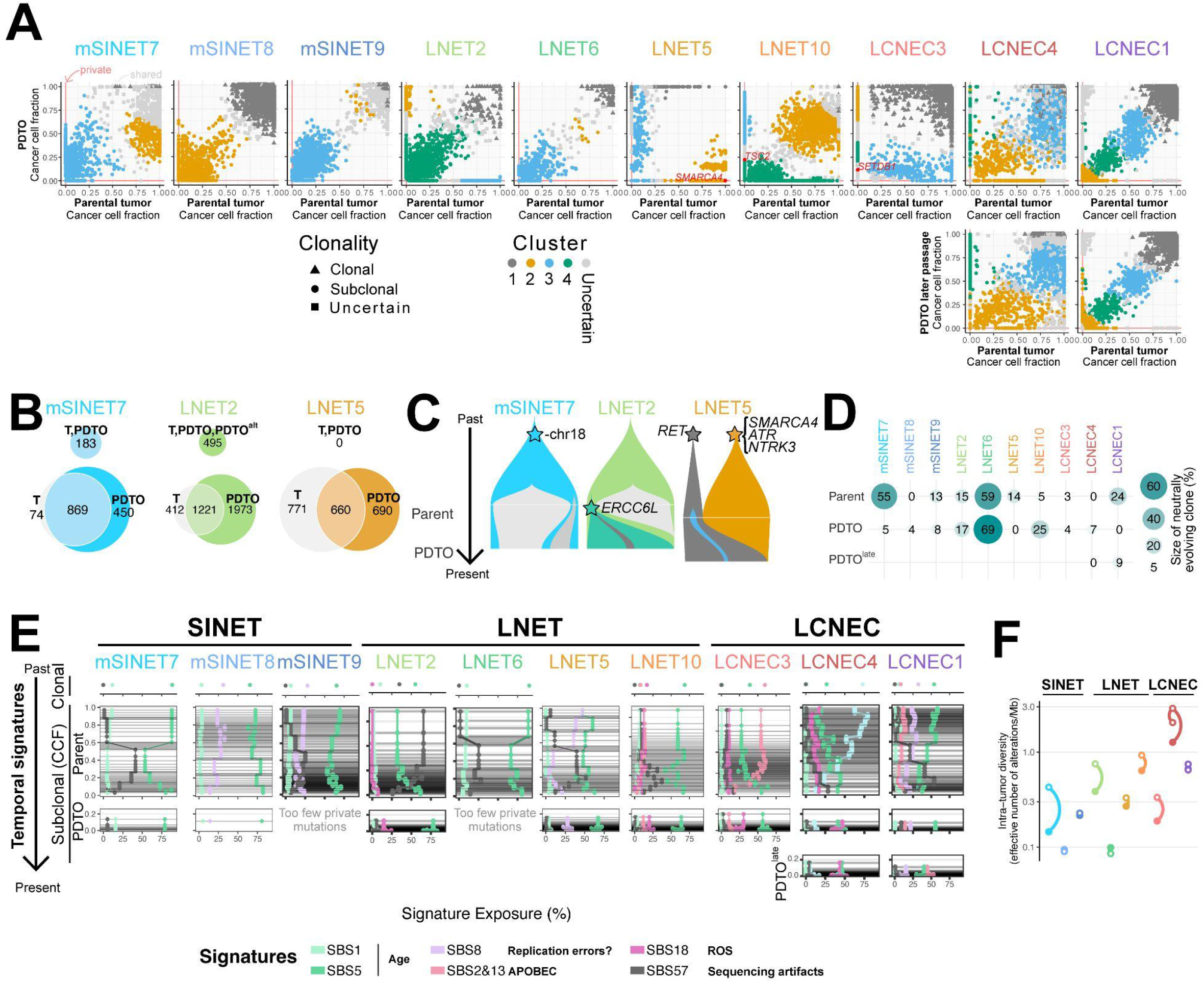
Extended Figure 5 including mixed PDTOs. (A) Clustering of joint cancer cell fractions (CCF) of small variants in regions with clonal copy number alterations. Colors correspond to clusters, and shapes to clonality. (B) Venn-Euler diagrams of shared and private clonal (top) and subclonal (bottom) somatic small variants from mixed samples. (C) Fish plots showing clonal reconstruction of tumor and organoid from mixed samples. (D) Extended Figure 5(C), including both high-purity and mixed samples. (E) Extended Figure 5(D), including both high-purity and mixed samples, and proportions of alterations from artifactual mutational signature SBS57. (F) Extended Figure 5(E): intra-tumor genetic diversity in both high-purity and mixed samples. See **Table S4**.

**Supplementary Figure 6.**
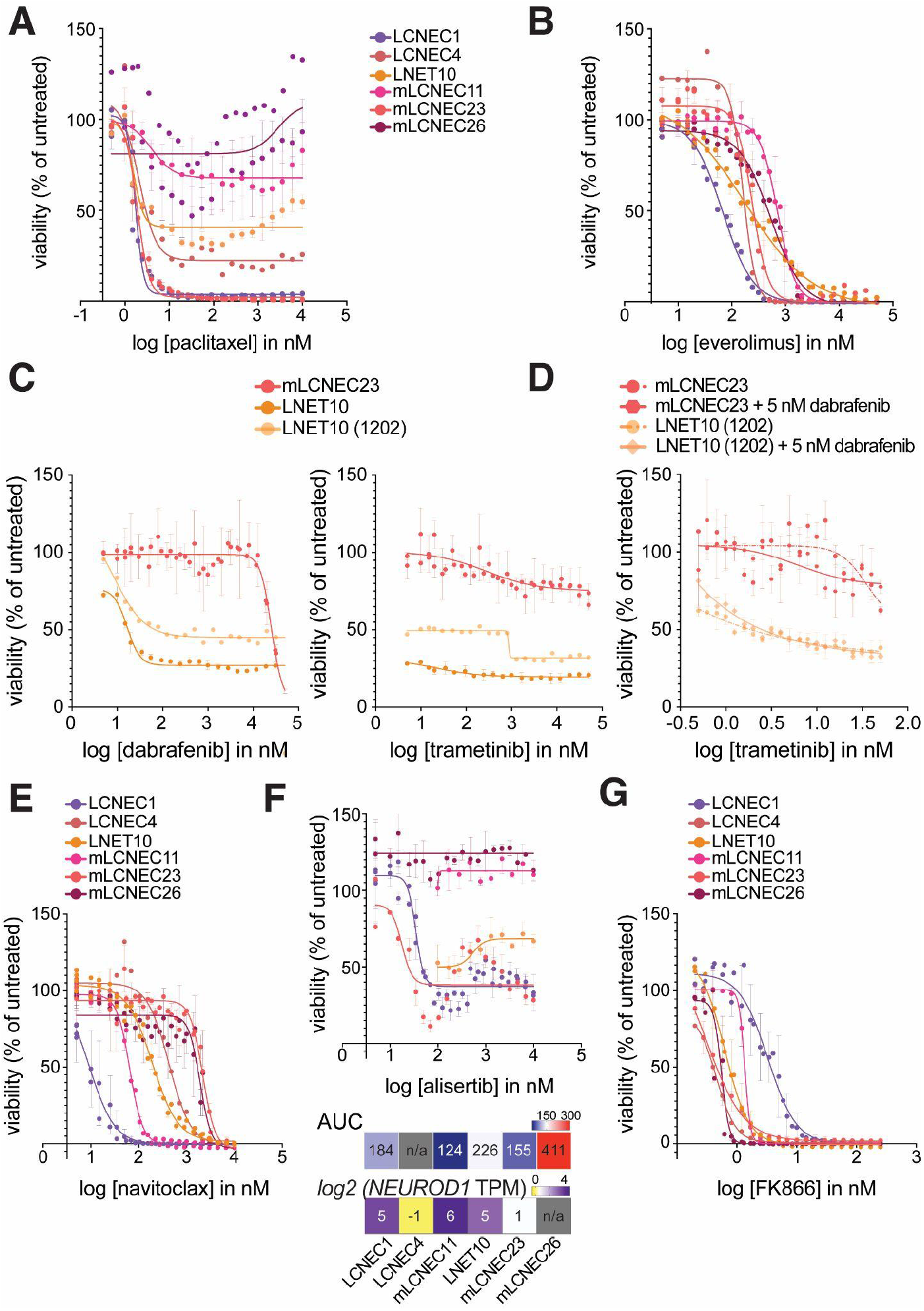
Dose-response curves for (A) Paclitaxel or (B) Navitoclax. Dots and error bars represent the mean and SEM from assays repeated on different days, respectively (*n* = 3-4), except for LCNEC4 where they represent technical replicates from one assay. Legend in A applied to B. (C) Dose-response curves for Dabrafenib or Trametinib. Dots and error bars represent the mean and SEM from assays repeated on different days (*n* = 3), respectively. Shown also are the values from an assay on LNET10, (LNET10 (1202)), which correspond to one technical replicate that deviated from the others. (D) Dose-response curves for treatments with Trametinib alone or in combination with 5nM Dabrafenib. Dots and error bars represent the mean and SEM from technical replicates (*n* = 3), respectively. (E) Dose-response curves for Navitoclax. Dots and error bars represent the mean and SEM from assays repeated on different days, respectively (*n* = 3-4), except for LCNEC4 where they represent technical replicates from one assay. Legend also applies to F and G. (F) Dose-response curves for Alisertib. Dots and error bars represent the mean and SEM from technical replicates (*n* = 3), respectively. Bottom panels show expression values for *NEUROD1*, and the AUC calculated for all samples tested. (G) Dose-response curves for FK866. Dots and error bars represent the mean and SEM from assays repeated on different days, respectively (*n* = 3-4), except for LCNEC4 and mLCNEC11 where they represent technical replicates from one assay.

**Supplementary Figure 7.**
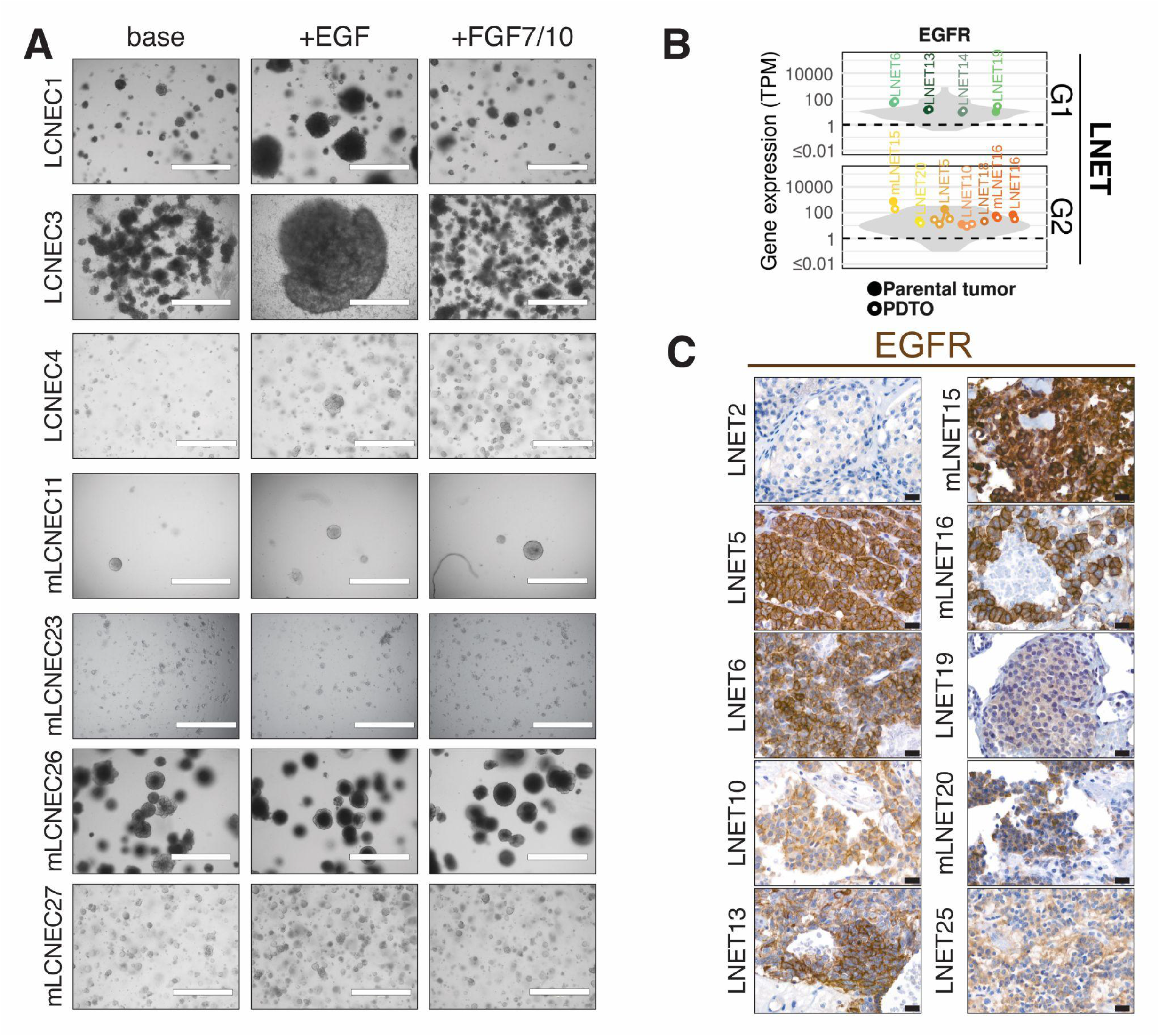
(A) Brightfield images showing LCNEC PDTO outgrowth in base NEN media and media supplemented with EGF or with FGF7 and FGF10. *Note LCNEC3 shows reduced cell adhesion in +EGF but growth rate is comparable to noEGF condition*. Scale bar: 1000 μm (B) Expression of EGFR parental tumors and matched PDTOs, in units of transcripts per million (TPM), for pulmonary NETs of different grades. Gray violin plots represent reference profiles with matching histological type and grade (*n*=75 for G1 LNET, *n*=40 for G2 LNET) (C) Immunohistochemical staining for the EGF receptor, EGFR, in parental tumor tissue for PDTO lines reported in this manuscript. Scale bar: 20 μm See **Table S2**.

Table S1. Clinical Information, passage times, and MKI67 expression of established PDTO lines, related to Figures 1 and 2

Table S2. Marker gene expression values, UMAP dimensions, PLS components, and PLS GSEA details, related to Figures 3 and S3

Table S3. PDTO purity classification details, related to Figures 3 and S3

Table S4. Mutations detected by WGS and RNAseq, related to Figures 4, S4, 5, and S5

Table S5. EGFR IHC details and scores, related to Figures 7 and S7

